# Small molecule inhibitors of transcriptional Cyclin Dependent Kinases impose HIV-1 latency, presenting “block and lock” treatment strategies

**DOI:** 10.1101/2023.08.17.553719

**Authors:** Riley M. Horvath, Zabrina L. Brumme, Ivan Sadowski

**Author notes:** Correspondence to: I. Sadowski, Dept. of Biochemistry and Molecular Biology, UBC, 2350 Health Sciences Mall, Vancouver, B.C., V6T 1Z3, CANADA.

## Abstract

Antiretroviral therapy is not a cure for HIV-1 as viral rebound ensues immediately following discontinuation. The block and lock therapeutic strategy seeks to enforce proviral latency and durably suppress viremic reemergence in the absence of antiretroviral therapy. Transcriptional Cyclin Dependent Kinase activity regulates LTR transcription, however, the effect and therapeutic potential of inhibiting these kinases for enforcing HIV-1 latency remains unrecognized. Using newly developed small molecule inhibitors that are highly selective for either CDK7 (YKL-5-124), CDK9 (LDC000067), or CDK8/19 (Senexin A), we found that targeting any one of these kinases prevented HIV-1 expression at concentrations that showed no toxicity. Furthermore, although CDK7 inhibition induced cell cycle arrest, inhibition of CDK9 and/or CDK8/19 did not. Of particular interest, proviral latency as induced by CDK8/19 inhibition was maintained following drug removal while CDK9 inhibitor induced latency rebounded within 24 hrs of discontinuation. Our results indicate that the Mediator complex kinases, CDK8/CDK19, are attractive block and lock targets while sole disruption of P-TEFb is unlikely to be efficacious.

## Introduction

The barrier towards a cure for HIV-1 is the population of extremely long-lived CD4^+^ T cells that harbor latent provirus. Antiretroviral therapy (ART) is capable of halting disease progression but has no effect on the latent HIV-1 reservoir and viral rebound proceeds immediately following discontinuation of treatment (Davey *et al*, 1999) (Zhang *et al*, 2000). The seemingly antithetical “shock and kill” and “block and lock” HIV-1 therapeutic strategies are currently being pursued to eliminate the current requirement of lifetime ART adherence (Sadowski & Hashemi, 2019). These aforementioned strategies rely on modulating proviral latency with “shock and kill” involving pharmacological reactivation mediated by latency reversal agents (LRAs) allowing for the subsequent destruction of infected cells through viral cytotoxicity and immunological surveillance (Kim *et al*, 2018) (Yeh & Ho, 2021). Unfortunately, formidable barriers to “shock and kill” strategies exist as it may be impossible to achieve reactivation of the majority of latent provirus (Ait-Ammar *et al*, 2020) (McMahon *et al*, 2021) (Siliciano & Siliciano, 2021) and it has become apparent that HIV-1 specific immune-mediated clearance is inefficient (Gunst *et al*, 2020) (Powell *et al*, 2020). Given the barriers to viral reactivation and clearance, the more feasible approach may be the alternative “block and lock” strategy by which durable latency is enforced to suppress viral rebound in the absence of ART (Moranguinho & Valente, 2020) (Vansant *et al*, 2020).

Cyclin Dependent Kinases (CDKs) are a family of proteins that catalyze the phosphorylation of serine/threonine residues, an activity that requires association with a cognate cyclin subunit. CDKs have historically been classified as either cell cycle associated kinases (including CDK1, CDK2, CDK4 and CDK6) or RNA polymerase II transcription associated kinases (CDK7, CDK8, CDK9, CDK12, CDK13 and CDK19). The multiprotein RNAPII complex mediates the synthesis of mRNA from a DNA template via a series of highly coordinated events that begins with the assembly of RNAPII and general transcription factors (GTFs) at a gene promoter. Initiation of RNA synthesis is coupled with Serine 5 (Ser5) and Serine 7 (Ser7) phosphorylation of the RNAPII Rpb1 subunit within the heptapeptide repeats of the carboxy-terminal domain (CTD) as mediated by the TFIIH associated kinase, CDK7 (Buratowski, 2009) (Rodríguez-Molina *et al*, 2016). Ser5 phosphorylation (Ser5-P) frees RNAPII from associated GTFs and the Mediator complex, facilitating promoter release (Rodríguez-Molina *et al*, 2016) (Wong *et al*, 2014). Additionally, CTD Ser5-P stabilizes the nascent transcript as the modification is recognized by enzymes that catalyze 7-methylguanosine (m7GTP) capping of the mRNA 5’ terminus (Ho & Shuman, 1999) (Fabrega *et al*, 2003) (Tome *et al*, 2018). In contrast, the effect of Serine 7 phosphorylation (Ser7-P) is ill-defined but is thought to prime subsequent CTD phosphorylation steps (Czudnochowski *et al*, 2012).

CDK7 initiates transcription from the core promoter, however, RNAPII may be associated with the pause factors DRB sensitivity-inducing factor (DSIF) and negative elongation factor (NELF) which cause stalling 20-60 nucleotides downstream of the transcriptional start site (TSS) in an effect known as promoter proximal pausing (Core & Adelman, 2019). Recruitment of the Cyclin T1/ CDK9 heterodimeric P-TEFb complex catalyzes RNAPII CTD Serine 2 phosphorylation (Ser2-P), a post-translational mark that is also deposited by CDK12 and CDK13 (Bartkowiak *et al*, 2010) (Blazek *et al*, 2011). In addition to Ser2-P, CDK9 phosphorylates DSIF and NELF causing the former to be converted to an elongation factor and the latter to be released, enabling processive elongation (Yamaguchi *et al*, 2013). As RNAPII reaches the 3’ end of the gene, the CTD is dephosphorylated, elongation transitions to a slowed pace, the mRNA is cleaved, and transcription is terminated (Cortazar *et al*, 2019) (Huang *et al*, 2020).

The 26-subunit Mediator complex facilitates genome-wide transcription by bridging sequence-specific DNA-binding transcription factors (TFs) with RNAPII. A quaternary complex of Cyclin C, CDK8 or CDK19, MED12, and MED13 forms the Mediator kinase module that associates reversibly with Mediator. Unlike the Mediator complex, the kinase module is not a genome-wide transcriptional co-activator. Rather, the Mediator kinase module selectively regulates transcription by phosphorylating transcription factors. The outcome of CDK8/19 kinase activity can be activation or repression depending on the functional effect of the modification.

For instance, CDK8/19 phosphorylation of β-catenin (Firestein *et al*, 2008), NFκB (Chen e*t al*, 2017) (Li e*t al*, 2019), SREBP-1 (Zhao *et al*, 2012), SMAD1/3 (Alarcón e*t al*, 2009), and STAT1 (Bancerek e*t al*, 2013) (Steinparzer e*t al*, 2019) enhances transcriptional activity while phosphorylation of Notch ICD promotes its degradation leading to target gene repression (Fryer *et al*, 2004) (Kuang e*t al*, 2020).

CDK7, CDK9, and CDK8/19 are transcriptional Cyclin Dependent Kinases (tCKDs) that are involved in HIV-1 expression. Recruitment of CDK7 (TFIIH) to the HIV-1 LTR has been described as the rate limiting step of latency reversal (Kim e*t al*, 2006) while the viral trans-activator Tat recruits CDK9 (P-TEFb) through interaction with nascent TAR RNA resulting in enhanced transcriptional elongation. Interestingly, the host cellular factor TRIM24 has been observed to stimulate RNAPII HIV-1 promoter clearance, an effect that is associated with enrichment of CDK9 and Ser2-P at the LTR (Horvath e*t al*, 2023). Moreover, the TRIM24 bromodomain inhibiter IACS-9571 and the BRD4 bromodomain inhibiter JQ1 reverse latency by promoting association of CDK9 with the LTR (Zhu *et al*, 2012) (Li *et al*, 2013) (Horvath e*t al*, 2023). Given the role of CDK9 for HIV-1 expression, the kinase represents a potential anti-viral target (Canduri *et al*, 2008) (de la Fuente *et al*, 2003). Inhibitors of CDK9 such as purine derivatives (Roscovitine) and flavonoids (flavopiridol) have been shown to repress HIV-1 replication (Wang e*t al*, 2001) (Schang *et al*, 2002) (Schonhofer e*t al*, 2021). However, these drugs lack specificity, targeting numerous CDKs (CDK2, CDK5, CDK7, etc.) in addition to other host cell enzymes (Meijer *et al*, 1997) (Schonhofer *et al*, 2021) that results in global inactivation of RNAPII transcription and cell cycle arrest (Blagosklonny, 2004) (Chao & Price, 2001). In addition to the aforementioned tCDKs, a role for CDK8 as a co-activator of HIV-1 transcription has recently been described with inhibitors of CDK8/19 causing repression of proviral expression both *in vitro* and e*x vivo* (Horvath e*t al*, 2023), though the ability of CKD8/19 inhibition to enforce robust latency has not been investigated.

Only recently have specific small molecule inhibitors of CDK7 and CDK9 been developed. Given this technological advancement, we sought to determine the HIV-1 “block and lock” therapeutic potential of targeting CKD7, CDK9, and/or CDK8/19. Treatment with YKL-5-124 (CDK7 inhibitor) (Olson *et al*, 2019), LDC000067 (CDK9 inhibitor) (Albert *et al*, 2014), or Senexin A (CDK8/19 inhibitor) (Porter *et al*, 2012) was able to prevent T cell activation mediated proviral induction in JLat Jurkat latency models and primary CD4^+^ peripheral blood monocytes (PBMCs) isolated from people living with HIV-1. Although each of the small molecules suppressed proviral expression, we observed synergistic inhibition when applied in combination with LDC000067 (CDK9i) and Senexin A (CDK8/19i) displaying the greatest magnitude of synergy. Although inhibition of CDK7 with YKL-5-124 was unable to affect proviral expression without causing cell cycle defects, CDK9 and CDK8/19 inhibition was extremely well tolerated over long durations of exposure. Given this tolerance, we were able to establish drug induced latency through multi-day treatment of HIV-1 infected cells with LDC000067 and/or Senexin A. Upon drug exclusion, cells treated with the CDK9 inhibitor LDC000067 displayed immediate viral rebound. In contrast, Senexin A induced latency was found to be robust, lasting following discontinuation. Collectively, our results suggest that the therapeutic potential of P-TEFb disruption is severely limited while CDK8/19 inhibitors are attractive latency promoting agents in the “block and lock” strategy to a functional HIV-1 cure.

## Results

### tCDK activity is required for HIV-1 induction in response to T cell activation

We sought to examine the requirement of tCDK activity for HIV-1 expression using the small molecule inhibitors YKL-5-124 (CDK7i), LDC000067 (CDK9i), and Senexin A (CDK8/19i) (Fig. 1*A*-*C*). These recently developed chemical inhibitors display vastly improved target selectivity over previously developed CDK inhibitors (Olson *et al*, 2019) (Albert *et al*, 2014) (Porter *et al*, 2012). Exposure of Jurkat human T cells to increasing concentrations of inhibitor determined half lethal concentration’s (LC50) of 93 μM, 499 μM, and 200 μM for YKL-5-124 (CDK7i), LDC000067 (CDK9i), and Senexin A (CDK8/19i), respectively (Fig. 1*D*). Having determined non-toxic concentration ranges, we investigated the effect of these drugs using the previously characterized JLat10.6 and JLat9.2 models of HIV-1 latency. JLat Jurkat human T cell lines possess a full-length HIV-1 provirus where GFP replaces *Nef*, and expression of the fluorescent protein is driven by the 5’ LTR (Jordan *et al*, 2003). Treatment with YKL-5-124, LDC000067, or Senexin A displayed dose-dependent suppression of HIV-1 induction in response to PMA/ionomycin mediated T cell activation in the JLat10.6 cell lines as measured by the percent of cells expressing GFP and the GFP Mean Fluorescent Intensity (MFI) (Fig. 2*A*-*C*). Here, YKL-5-124 (CDK7i) displayed the most potent activity with a half inhibitory concentration (IC50) for cellular GFP expression of 0.11 μM while LDC000067 and Senexin A followed with GFP expression IC50’s of 1.42 μM and 15.43 μM, respectively (Fig. 2*B*). Comparable results were obtained when GFP MFI was assessed (Fig. 2*C*). Furthermore, a similar inhibitory effect was observed for the JLat9.2 latency model (Fig. 2*D*), with lower concentrations of inhibitor being required to repress proviral expression as compared to the JLat10.6 cell line (Fig. 2*E*, 2*F*). YKL-5-124 was again the most potent inhibitor (IC50: 0.02 μM), followed by Senexin A (IC50: 0.43 μM) and LDC000067 (IC50: 0.56 μM). Notably, cellular viability of JLat10.6 and JLat9.2 cellse were not affected by tCDK inhibition (Fig. S1).

**Figure 1.**
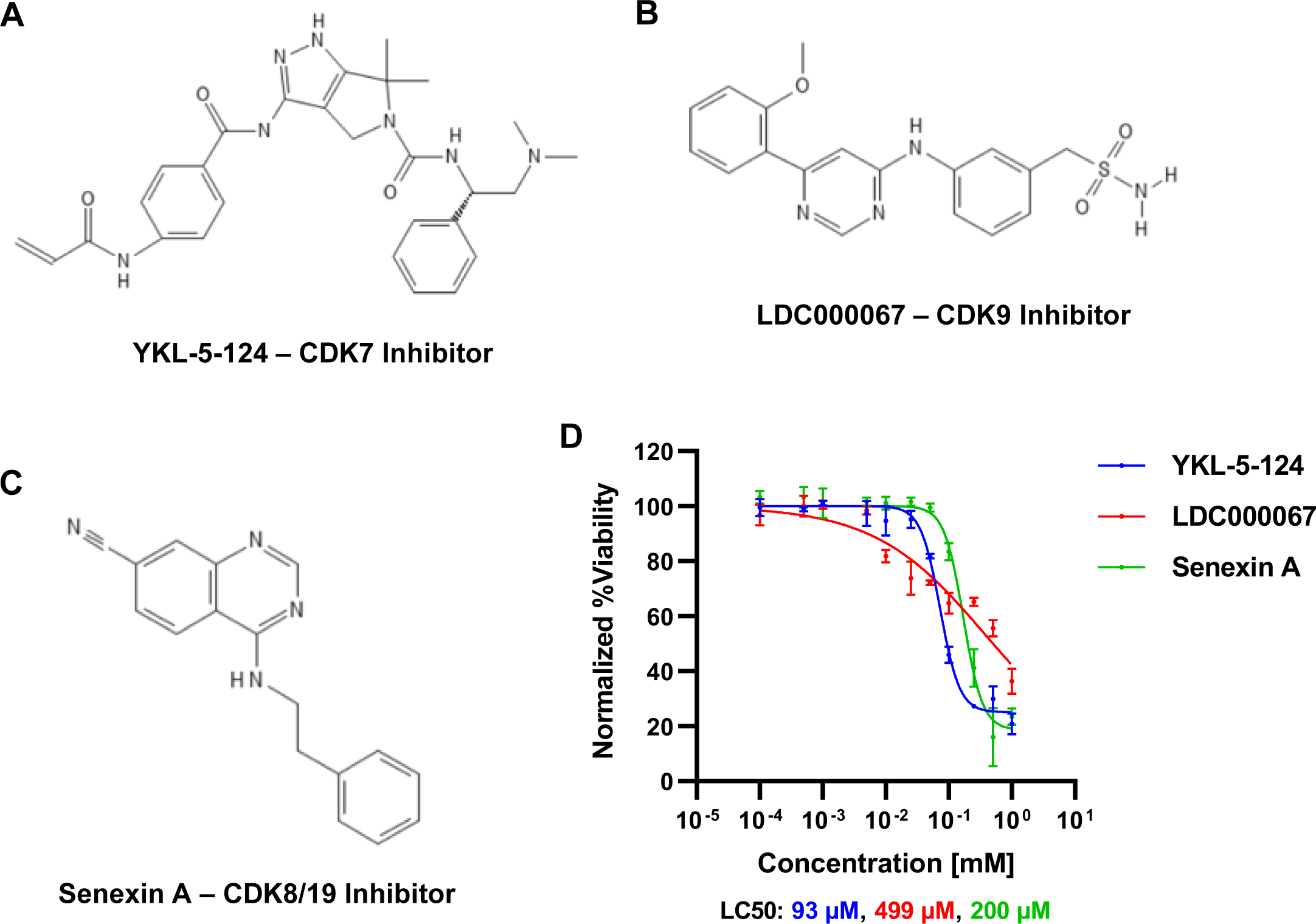
Chemical inhibition of tCDKs is well tolerated in T cells. **A-C:** Molecular structure of YKL-5-124 (CDK7 inhibitor) (**A**), LDC000067 (CDK9 inhibitor) (**B**), and Senexin A (CDK8 and CDK19 inhibitor) (**C**). **D:** Jurkat human T cells were treated with increasing concentrations of the indicated compound for 24 hrs after which cellular viability was determined. Results are normalized to untreated control (*n* = 2, mean ±LSD).

**Figure 2.**
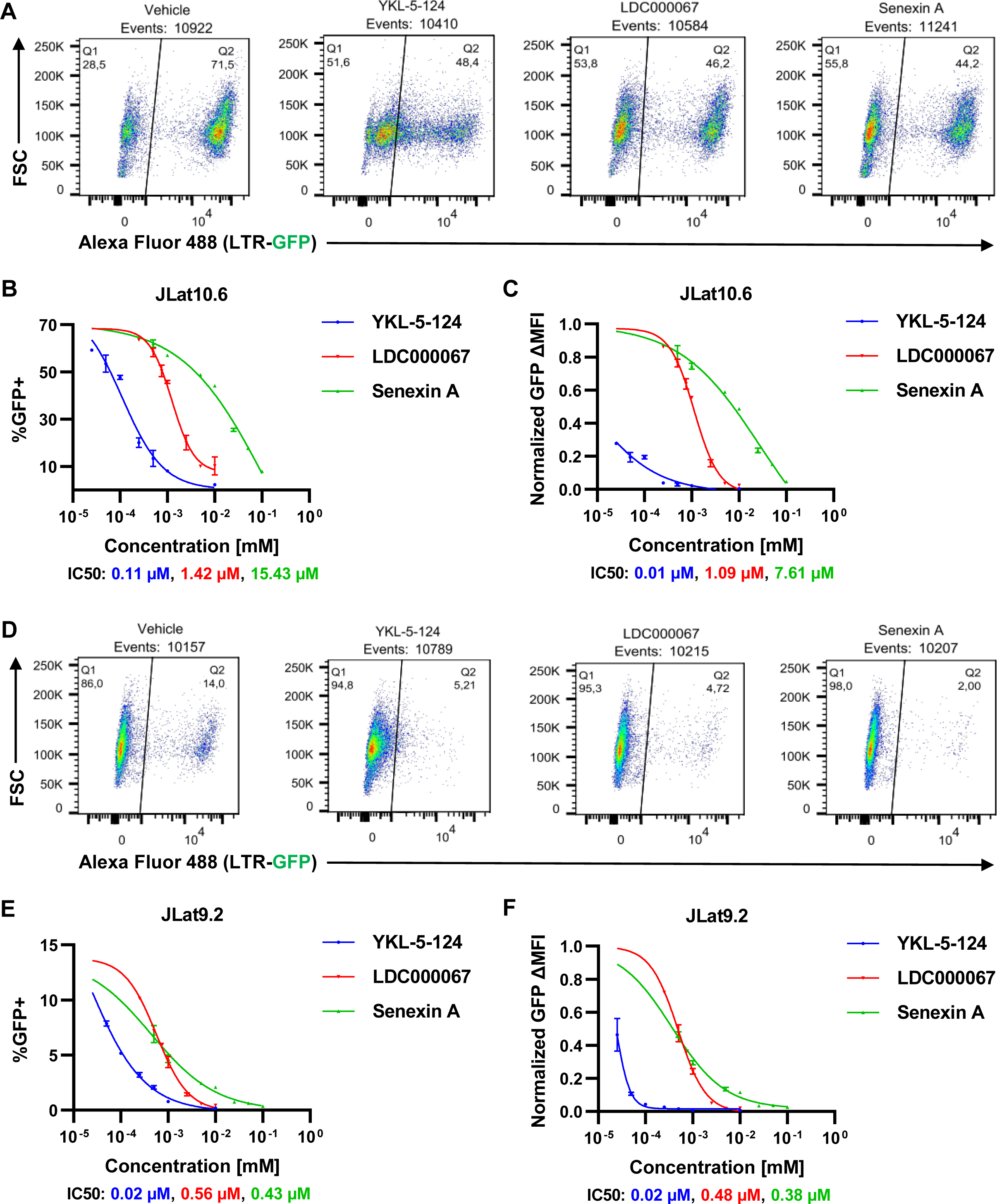
HIV-1 latency cell line models require tCDK kinase activity for reactivation. **A:** Representative flow cytometry scatter plots of JLat10.6 cells that were pre-treated for 30 min with DMSO (Vehicle), 100 nM YKL-5-124 (CDK7i), 1 μM LDC000067 (CDK9i), or 10 μM Senexin A (CDK8/19i) and subsequently incubated with 5 nM PMA and 1 μM ionomycin for 20 hrs. **B, C:** Following 30 min pre-treatment with the indicated concentration of tCDK inhibitor, JLat10.6 cells were incubated with 5 nM PMA and 1 μM ionomycin. After 20 hrs, flow cytometric analysis was performed to determine HIV-1 expression as reported as the proportion of GFP positive cells (**B**) and the GFP delta (Δ) Mean Fluorescent Intensity (MFI) (MFI of treatment subtracted by the MFI of untreated) (**C**) (*n* = 2, mean ±LSD). **D:** As in (**A**), but JLat9.2 cells were assessed. **E, F:** As in B, C, but JLat9.2 cells were examined (*n* = 2, mean ±LSD).

The JLat10.6 and JLat9.2 cell lines display a distinct pattern of proviral activation in response to T cell activation, whereby cells tend to possess provirus that are either inactive or fully expressed (Fig. 2*A*, 2*D*, Vehicle). Treatment with LDC000067 (CDK9i) or Senexin A (CDK8/19i) diminished the proportion of GFP+ cells (Fig. 2*B*, 2*E*), however, the bimodal expression pattern remains apparent where there is still the tendency for those provirus that escape latency to be fully active (Fig. 2*A*, 2*D*, LDC000067, Senexin A). In contrast, treatment with the CDK7 inhibitor YKL-5-124 does not follow bimodal expression. Rather, a gradient of expression is produced with many proviruses displaying activation levels that verge on the GFP-/GFP+ border (Fig. 2*A*, 2*D*, YKL-5-124). The viral encoded trans-activator of transcription Tat protein produces a strong positive feedback loop. CDK7 is involved in initiation of transcription, ostensibly a Tat independent process, while CDK9 is recruited by Tat to the LTR to facilitate release of RNAPII promoter proximal pausing. As such, it is probable that the different expression pattern observed between YKL-5-124 inhibition of CDK7 and either LDC000067 inhibition of CDK9 or Senexin A inhibition of CDK8/19 is the result of Tat independent versus Tat dependent transcriptional induction.

### tCDK inhibitors synergistically inhibit HIV-1 expression

CDK7, CDK9, and CDK8/19 regulate different stages in RNAPII gene expression. As such, it would be expected that combined treatment with inhibitors specific for particular tCDK’s would reveal synergistic interaction. To examine synergistic inhibition, we treated JLat10.6 cells with YKL-5-124 (CDK7i) and LDC000067 (CDK9i) (Fig. 3*A*), YKL-5-124 (CDK7i) and Senxin A (CDK8/19i) (Fig. 3*B*), or LDC000067 (CDK9i) and Senexin A (CDK8/19i) (Fig. 3*C*), and induced proviral expression with the addition of PMA/ionomycin. Co-treatment with any pair of tCDK inhibitor was found to have a greater repressive effect than mono-treatment as indicated by the percent of GFP positive cells following reactivation (Fig. 3*A*-*C*). Additionally, cellular viability was not affected by any of the applied treatments (Fig. S2). We confirmed synergistic inhibition by performing Bliss independence modelling (Laird *et al*, 2014) (Liu *et al*, 2018) and found that the combination of LDC000067 (CDK9i) and Senexin A (CDK8/19) displayed the strongest interaction (Fig. 3*D*-*F*). Of note, the apparent synergy dips at higher concentrations of chemical inhibitor as a result of viral expression approaching zero (Fig. 3*D*-*F*). These observations are consistent with the tCDK inhibitors being reported as selective for either CDK7, CDK9, or CDK8/19 and indicate that these kinases function through independent mechanisms.

**Figure 3.**
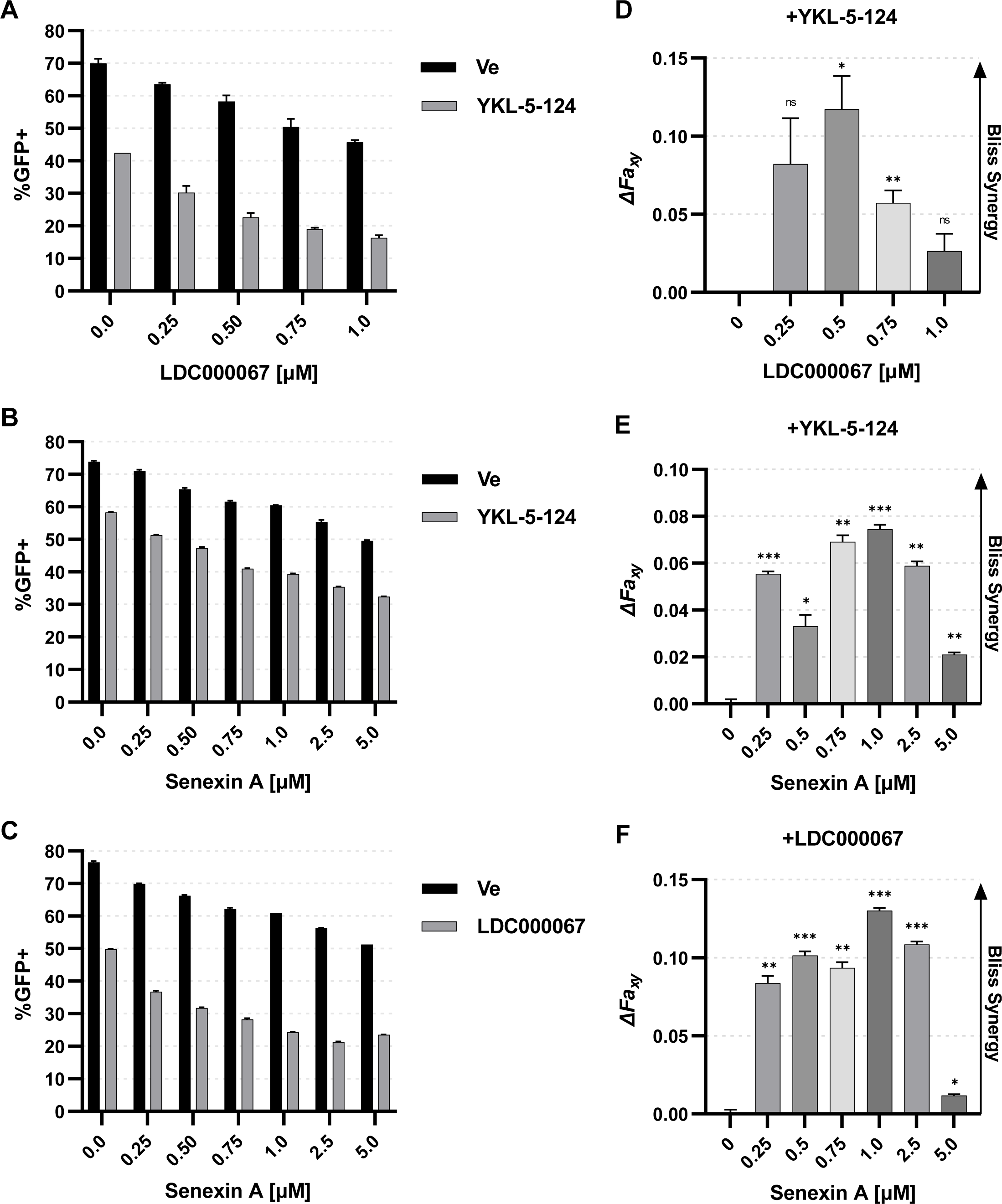
tCDK inhibitors synergistically inhibit HIV-1 transcription. **A - C:** JLat10.6 cells were pre-treated for 30 min with 100 nM YKL-5-124 (CDK7i) and the indicated concentration of LDC000067 (CDK9i) (**A**) or Senxexin A (CDK8/19i) (**B**), or were pre-treated 30 min with 1 μM LDC000067 and the indicated concentration of Senexin A (**C**). Subsequently, cells were incubated with 5 nM PMA/ 1 μM ionomycin for 20 hrs and analyzed by flow cytometry following 20 hrs (*n* = 2, mean ±LSD). **D - F:** Calculation of synergy between the indicated treatments in **A - C** using Bliss Independence Modeling. Data are presented as the difference between the predicted and the observed fractional HIV-1 expression response to the given drug combination (*n* = 2, mean ±LSD). See Materials and Methods for more details.

### Inhibition of tCDK activity antagonizes proviral reactivation in response to mechanistically diverse latency reversal agents

Having observed that tCDK inhibition suppresses reactivation of latent provirus in response to T cell activation, we next examined the effect on a variety of mechanistically diverse latency reversal agents (LRAs). As mentioned above, co-treatment with the phorbol ester PMA and the ionophore ionomycin triggers T cell activation by stimulating Ras-MAPK, PKC-NFκB, and calcineurin-NFAT. Ingenol 3-angelate (PEP005) activates PKC-NFκB, partially mimicking T cell activation (Kedei *et al*, 2004). JQ1 and IACS-9571 are bromodomain inhibitors that reverse latency as mediated by their respective targets, BRD4 (Li *et* al, 2013) and TRIM24 (Horvath *et al*, 2023). Finally, suberanilohydroxamic acid (SAHA) is a well-studied histone deacetylase inhibitor (HDACi) known to reactivate HIV-1 expression *in vitro* and *in vivo* (Contreras *et al*, 2009) (Desimio *et al*, 2017). Of the LRAs applied, the bromodomain inhibitors JQ1 and IACS-9571 had the smallest effect on proviral reactivation in the JLat10.6 latency model (Fig. 4*A*, 4*B*). The NFκB agonist PEP005 and the HDACi SAHA produced intermediate reactivation while co-treatment with either PEP005/IACS-9571 or PMA/ionomycin exhibited the greatest levels of HIV-1 expression (Fig. 4*A*, 4*B*).

**Figure 4.**
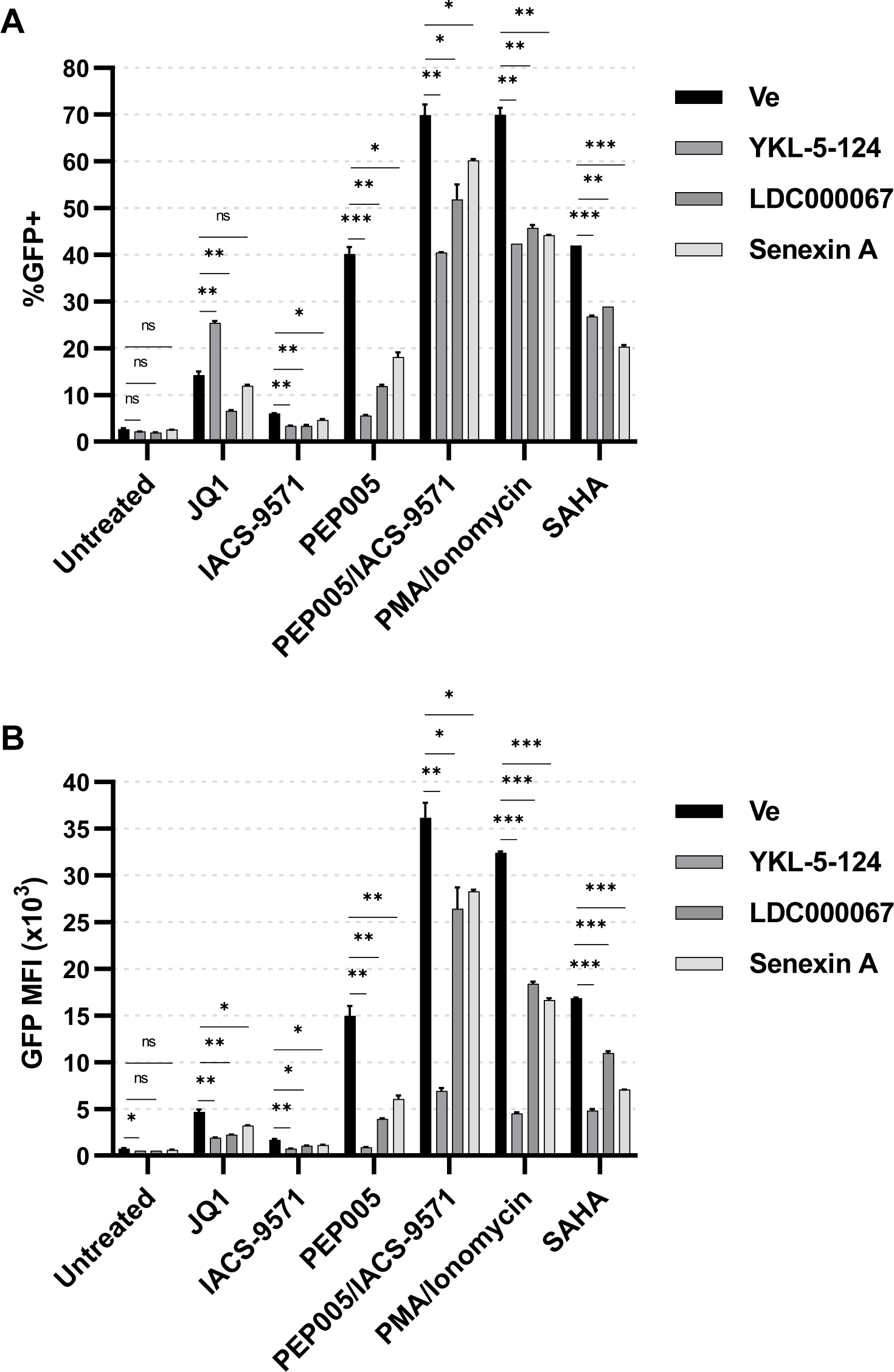
tCDK kinase activity mediates proviral expression in response to latency reversal agents. **A, B:** JLat10.6 cells were pre-treated for 30 min with 100 nM YKL-5-124 (CDK7i), 1 μM LDC000067 (CDK9i), 10 μM Senexin A (CDK8/19i), or a vehicle control (Ve, DMSO). Subsequently, cells were incubated with10 μM JQ1, 20 μM IACS-9571, 5 nM PEP005, 5 nM PEP005/ 20 μM IACS-9571, 5 nM PMA/ 1 μM ionomycin, 5 μM SAHA, or were left untreated. Following 20 hrs, viral expression was assessed by flow cytometry and reported as the percent of GFP positive cells (**A**) and the Mean Fluorescent Intensity (MFI) of GFP (**B**) (*n* = 2, mean ±LSD).

We applied 100 nM YKL-5-124 (CDK7i), 1 μM LDC000067 (CDK9i), or 10 μM Senexin A (CDK8/19i), concentrations near the inhibitor’s LC50 for PMA/ionomycin activation, in combination with the LRAs discussed above. CDK7 inhibition by YKL-5-124 limited proviral reactivation as measured by the population of GFP+ cells in response to every LRA other than JQ1 where oddly, we observe an increased proportion of GFP expressing cells (Fig. 4*A*). Interestingly, although CDK7 inhibition with concomitant JQ1 treatment produced a greater proportion of GFP+ cells (Fig. 4*A*), the GFP MFI was decreased as compared to the vehicle control (Fig. 4*B*) indicating that more cells expressed GFP but at attenuated levels upon introduction of YKL-5-124. Treatment with LDC000067 (CDK9i) or Senexin A (CDK8/19i) was found to stunt HIV-1 transcriptional induction in response to all of the tested LRAs (Fig. 4*A*, 4*B*). In agreement with previous reports (Horvath *et al*, 2023), we observed a vigorous synergistic interaction between PEP005 activation of NFκB and IACS-9571 inhibition of the TRIM24 bromodomain (Fig. S3). Moreover, tCDK inhibition was least effective at repressing the PEP005/IACS-9571 reactivation protocol, further demonstrating its latency reversal potential (Fig. 4*A*, 4*B*). Finally, although these concentrations of tCDK inhibitors equally suppressed the emergence of GFP positive cells in response to PMA/ionomycin T cell activation (Fig. 4*A*, PMA/Ionomycin), we found certain inhibitors to be more effective at combatting particular LRAs. For instance, Senexin A (CDK8/19i) was most effective at suppressing SAHA mediated activation (Fig. 4*A*, SAHA) while LDC000067 (CDK9i) was most potent at inhibiting expression in response to PEP005 (Fig. 4*A*). Taken together, these results show that tCDK inhibition suppresses proviral activation in response to mechanistically diverse LRAs while suggesting that the kinases have divergent functions for HIV-1 transcription.

### Tat dependence of tCDKs for activation of HIV-1 transcription

Next, we examined the Tat dependence of CDK7, CDK9, and CDK8/19 for activation of HIV-1 transcription. The Jurkat derived JLatA72 cell line possesses an integrated HIV-1 LTR that drives expression of GFP but does not encode Tat. Untreated JLatA72 cells are ∼30% GFP positive and this proportion increases to ∼60% upon PMA/ionomycin stimulation (Fig. 5*A*, 5*B*, Untreated, Vehicle). To examine the Tat dependence of the tCDKs, we treated JLatA72 with a kinase inhibitor and subsequently induced T cell activation with PMA/ionomycin. Surprisingly, we found that YKL-5-124 inhibition of CDK7 caused an increase in the proportion of GFP positive cells (Fig. 5*A*, 5*B*) while also lowering overall proviral expression (Fig. 5*C*), an effect that is strikingly similar to co-treatment with JQ1 (Fig. 4). The CDK9 inhibitor LDC000067 had no effect on the proportion of GFP positive cells (Fig. 5*A*, 5*B*) and only slightly decreased overall GFP expression (Fig. 5*C*), consistent with the role of Tat for recruitment of CDK9 to the HIV-1 LTR. Finally, inhibition of CDK8/19 with Senexin A decreased the emergence of GFP (Fig. 5*A*, 5*B*) while also suppressing overall GFP expression (Fig. 5*C*), indicating Tat independent regulation. We confirmed Tat dependence by additional examination of the CEM derived GXR-5 cell line. As with the JLatA72 cell line, The GXR-5 cells possess a chromosomally integrated HIV-1 LTR that drives GFP expression but does not express Tat. Unlike JLatA72 cells, GXR-5 cells are almost entirely GFP positive (Fig. S4*A*, Fig. S4*B*) though an increase in GFP expression following PMA/ionomycin treatment is observable by assessing GFP Mean Fluorescent Intensity (MFI) (Fig. S4*C*). Similar to the JLatA72 cell line, GXR-5 cells display severely attenuated GFP induction upon treatment with YKL-5-124 (CDK7i) or Senexin A (CDK/19i) (Fig. S4*C*). In contrast, LDC000067 (CDK9i) caused small but significant inhibition of GFP expression (Fig. S4*C*).

**Figure 5.**
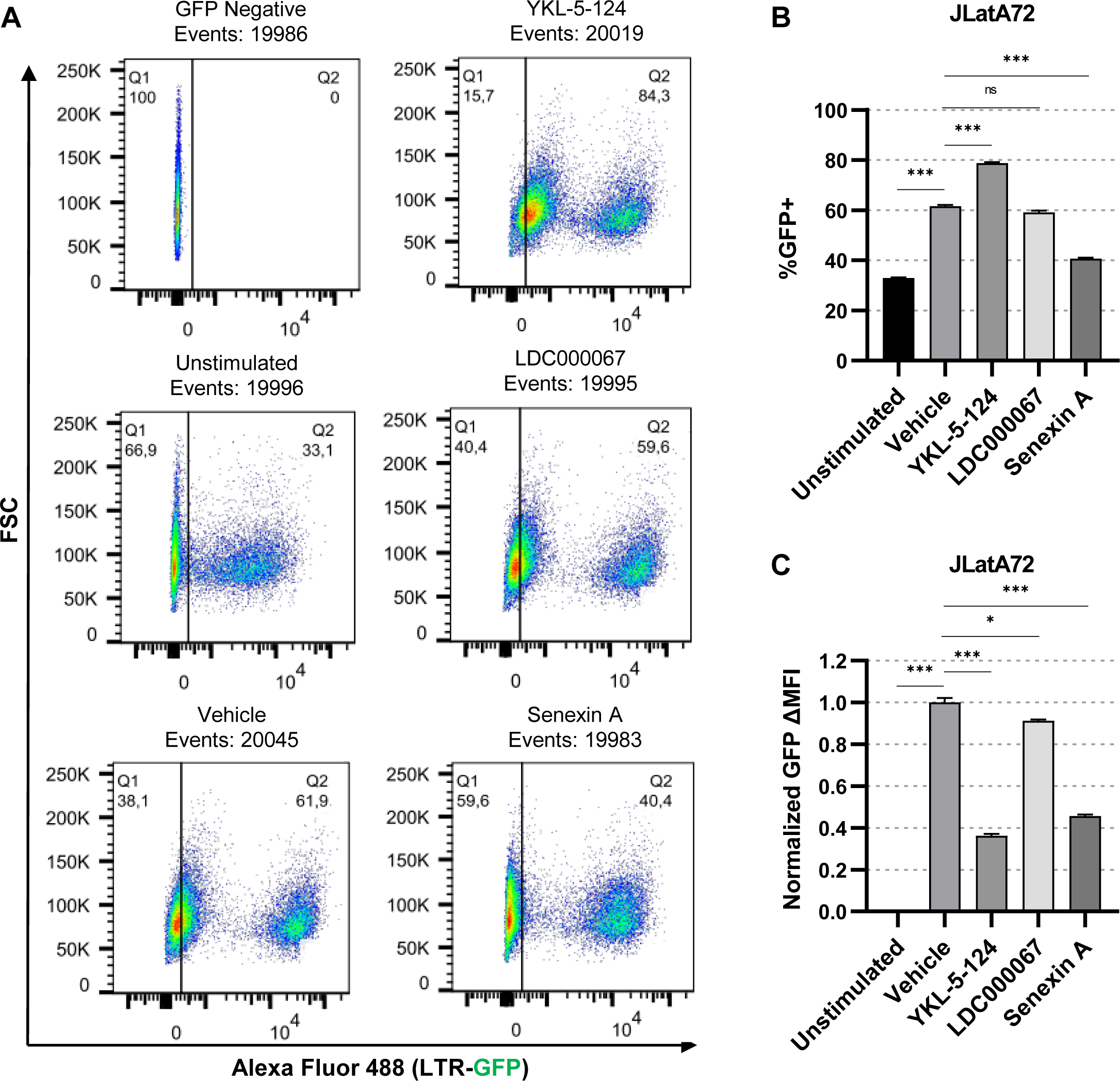
Tat dependency of tCDK kinase activity. **A:** Representative flow cytometry scatter plots of JLatA72 Jurkat cells (-Tat) that were pre-treated for 30 min with a vehicle control (DMSO), 100 nM YKL-5-124 (CDK7i), 1 μM LDC000067 (CDK9i), or 50 μM Senexin A (CDK8/19i) and subsequently incubated with 5 nM PMA and 1 μM ionomycin for 20 hrs. **B, C:** JLatA72 cells were left untreated or pre-treated for 30 min with a vehicle control (DMSO), 100 nM YKL-5-124, 1 μM LDC00067, or 50 μM Senexin A prior to being incubated with 5 nM PMA/ 1 μM ionomycin. Following 20 hrs, flow cytometry was performed with viral expression reported as the percent of GFP positive cells (**B**) and the delta GFP Mean Fluorescent Intensity (MFI) (**C**) (*n* = 2, mean ±LSD).

### Effect of tCDK inhibition on cell growth

Above, we examined the effects of acute tCDK inhibition on proviral expression. Given the stochastic nature of HIV-1 reactivation, a “block and lock” treatment strategy will likely necessitate longer periods of exposure to generate latency that persists subsequent to drug withdrawal. To further examine the cellular impacts of tCDK inhibition, we assessed the growth and viability of Jurkat human T cells exposed to a range of drug concentrations for 4-days. We found that inhibition of CDK7 with YKL-5-124 caused cell cycle arrest (Fig. 6*A*) at concentrations shown to inhibit HIV-1 expression (Fig. 2), a result that is consistent with CDK7 being the CDK-activating kinase (CAK) which phosphorylates and activates CDKs that have essential roles for cell-cycle progression (Fisher, 2005). Moreover, cellular toxicity begins to become apparent following 3- and 4-days of CDK7 inhibition (Fig. 6*B*). In contrast of CDK7 inhibition, inhibition of CDK9 with LDC000067 had no effect on cell division (Fig. 6*C*) or viability (Fig. 6*D*) at concentrations effective at limiting proviral expression (Fig. 2). Likewise, Senexin A inhibition of CDK8/19 had minimal impact on cell growth (Fig. 6*E*) and did not affect viability (Fig. 6*F*) within a range of concentrations that suppressed HIV-1 reactivation (Fig. 2). As CDK7 inhibition causes cell-cycle arrest at low concentrations, the therapeutic potential of YKL-5-124 and other CDK7 inhibitors will likely be limited. However, our results find that inhibition of CDK9 with LDC000067 or inhibition of CDK8/19 with Senexin is well tolerated over prolonged application within an optimal concentration range for HIV-1 repression.

**Figure 6.**
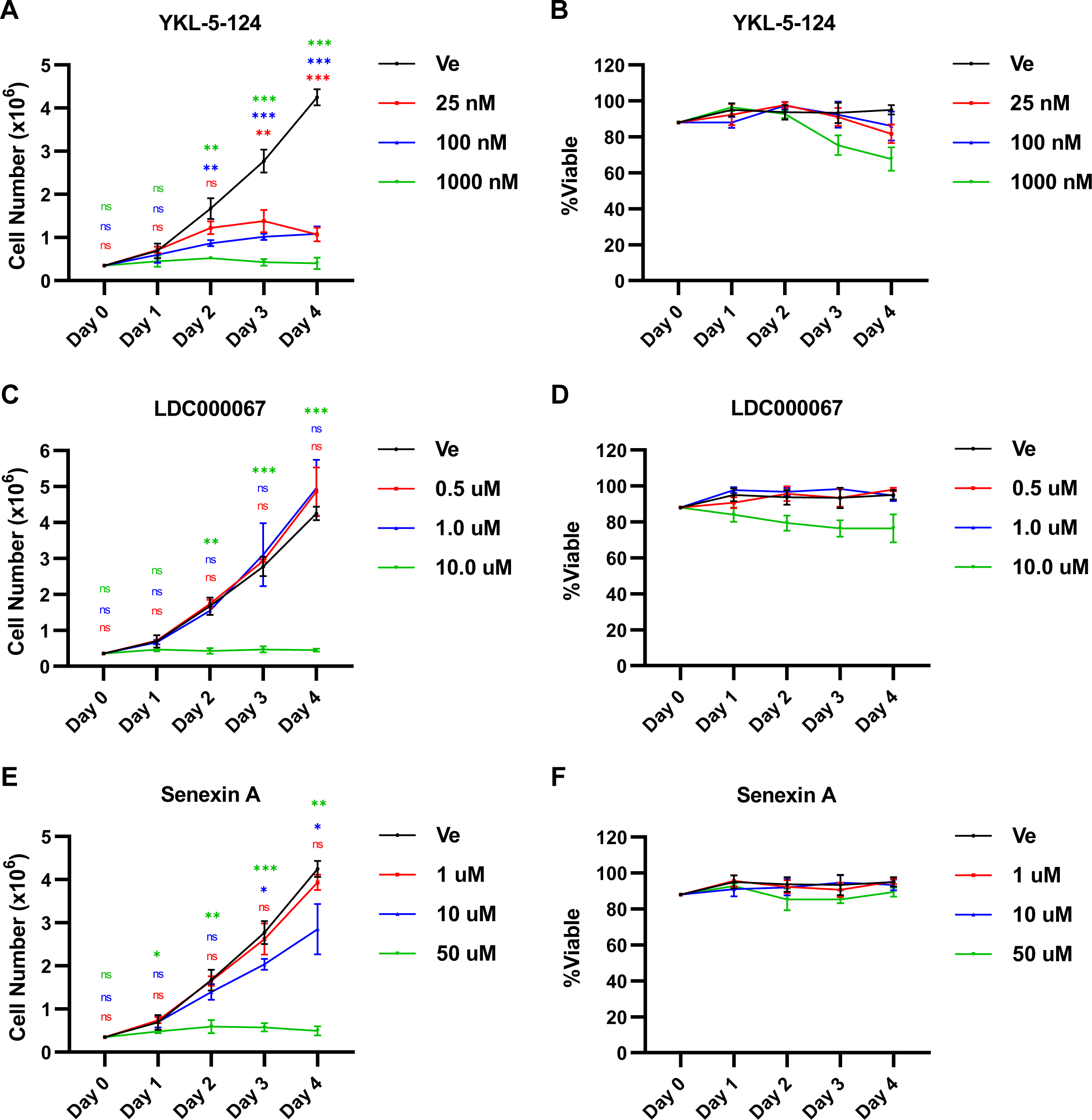
Effect of sustained tCDK inhibition on cellular growth and viability. **A, B:** Jurkat T cells were incubated with the indicated concentration of YKL-5-124 or a vehicle control (Ve, DMSO) with cell number (**A**) and viability (**B**) assessed on the indicated day. Media was replaced daily with inhibitor (*n* = 3, mean ±LSD). **C, D:** As in (**A**) and (**B**), but cells were treated with LDC000067 (*n* = 3, mean ±LSD). **E, F:** Performed as previously described but Jurkat cells were incubated with the indicated concentration of Senexin A (*n* = 3, mean ±LSD).

### Role of tCDKs for the establishment of immediate latency

Upon infection, nearly 50% of provirus are transcriptionally inactive, a phenomenon known as immediate latency (Dahabieh *et al*, 2013) (Dahabieh *et al*, 2014) (Battivelli e*t al*, 2018). Having observed that the tCDKs are required for reactivation of HIV-1 in response to various LRAs, we examined the effect of kinase inhibition on the occurrence of immediate latency using the Red-Green-HIV-1 (RGH) dual-reporter virus (Fig. 7*A*) (Dahabieh *et al*, 2013) (Dahabieh *et al*, 2014). RGH is a full-length HIV-1 construct that is rendered replication incompetent by mutation to *Env* and whereby eGFP expression is driven by the 5’ LTR and a constitutive PGK-mCherry cassette is inserted in place of *Nef*, allowing for differentiation between latent and productive proviral infection (Fig. 7*A*). To assess tCDK activity on immediate latency, we pre-treated Jurkat human CD4^+^ T cells with a kinase inhibitor and subsequently infected with RGH at a low multiplicity of infection (MOI). One day following infection, we either washed (removed) or maintained the drug inhibitor, assessing proviral latency 4-days post-infection by flow cytometry (Fig. 7*B*) (Fig. 7*C*). At the point of infection, inhibition of either CDK7 with YKL-5-124 or CDK8/19 with Senexin A was able to reduce the proportion of productive infections (Fig. 7*D*, Washed). Interestingly, inhibition of CDK9 with LDC000067 at the point of infection had no impact on immediate latency (Fig. 7*D*, Washed), suggesting that CDK9 and Tat are not involved in viral transcription early upon infection. In agreement with tCDK inhibition preventing reactivation of latent provirus (Fig. 2, Fig. 4), latency was enforced by all of the small molecule inhibitors when they were maintained for the 4 days (Fig. 7*D*, Maintained). Surprisingly, we found that YKL-5-124 inhibition of CDK7 caused the rate of infection to roughly double indicating that loss of CDK7 activity renders T cells susceptible to infection (Fig. 7*E*). Notably, although CDK7 inhibition causes an increase in HIV-1 infection (Fig 7*E*), integrants were enriched for latent phenotype (Fig. 7*C*, compare Untreated to YKL-5-124). These results indicate that CDK9 and CDK8/19 are attractive targets to suppress proviral expression.

**Figure 7.**
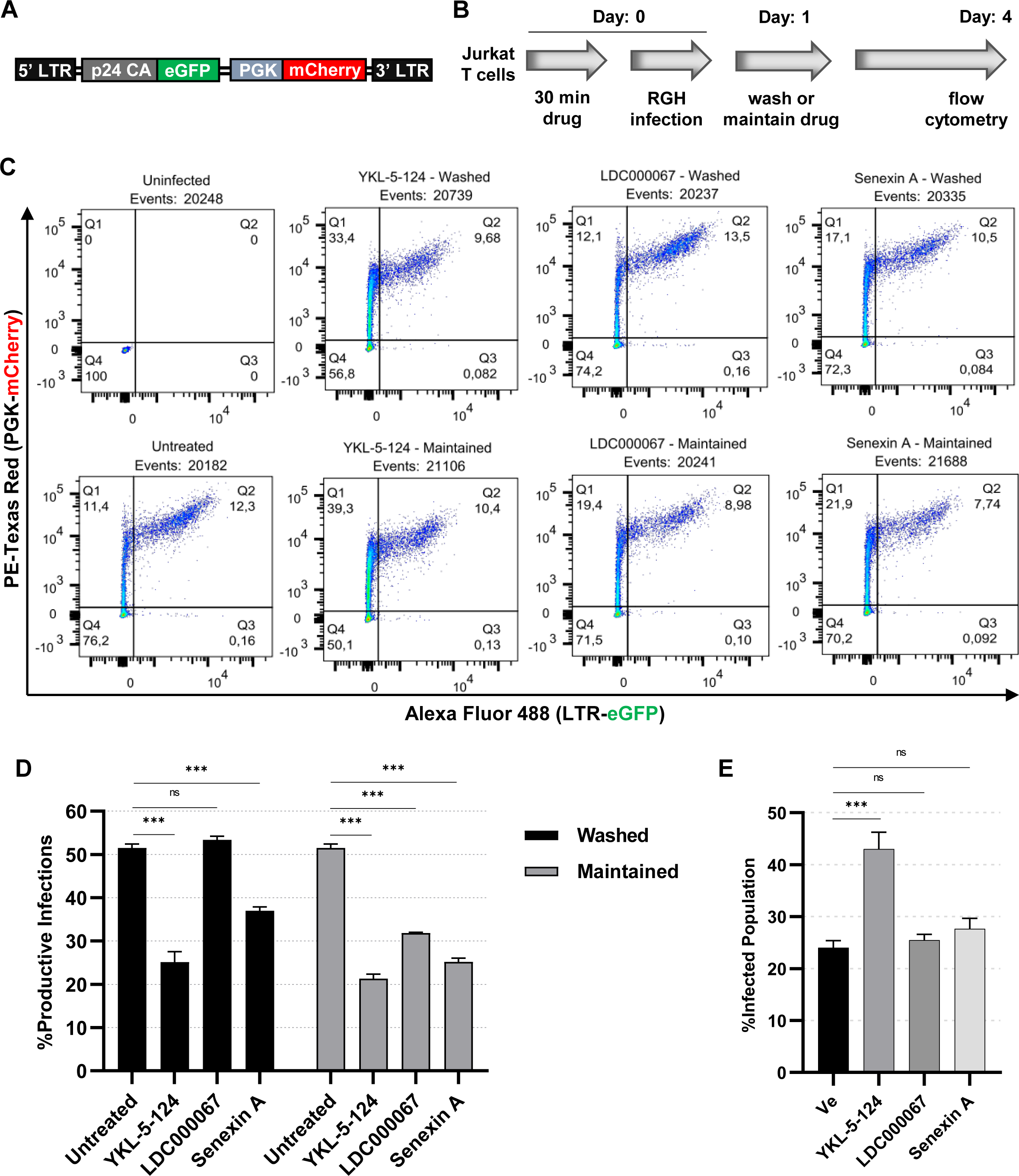
tCDK activity mediates the establishment of latency upon infection. **A:** Simplified depiction of the replication incompetent Red-Green-HIV-1 (RGH) dual reporter virus (Dahabieh *et al*, 2013). RGH is a full-length virus that is rendered to a single round of infection through mutation in *Env*, and that expresses GFP as driven by 5’ LTR transcription while also expressing mCherry from a constitutive PGK promoter that replaces *Nef,* allowing for latent and productive infections to be differentiated. **B:** Schematic representation of the RGH assay. Jurkat CD4^+^ human T cells were left untreated or were pre-treated for 30 min with 100 nM YKL-5-124, 1 μM LDC000067, or 10 μM Senexin A. Subsequently, cells were infected with RGH using equivalent amounts of virion. 1-day post-infection, media was refreshed and cells were either washed of the drug or maintained with the same concentration of inhibitor as prior. 4 days post-infection, proviral expression was assessed by flow cytometry. **C:** Representative flow cytometry scatter plots of RGH infected cells, treated as indicated in (**B**) and assessed 4 days post-infection. **D:** Jurkat T cells were treated as in (**B**), with the percentage of productive infections determined by flow cytometry 4 days post-infection. The %Productive Infections was determined from the ratio of GFP+/mCherry+ cells to all mCherry+ cells (*n* = 3, mean ±LSD). **E:** Bar graph summary of the infection rate for the indicated treatment determined by cells displaying PGK driven mCherry expression (*n* = 3, mean ±LSD).

### Disruption of CDK8/19 but not CDK9 enforces robust latency

An optimal “block and lock” therapy would involve generating proviral latency that is maintained subsequent to the removal of the latency promoting agent (LPA). To examine the capability of tCDK inhibitors to create robust and lasting latency, we applied LDC000067 (CDK9i) and/or Senexin A (CDK8/19i) to the previously described mdHIV #77 cell line (Hashemi *et al*, 2016). mdHIV #77 is a Jurkat Tat derived clonal line that contains a mini-dual reporter HIV-1 provirus (Fig. 8*A*) that is integrated into an intron of the *PACS1* gene in the (+) orientation (Hashemi *et al*, 2016). Although every cell of this clonal line contains the mini-virus integrated in the same location, stochastic bursts of transcription produce a slightly fluctuating population of ∼3% productive infections as indicated by the presence of LTR expressed dsRed and eGFP transcribed by an internal EF1α promoter (Hashemi *et al*, 2016). We applied the tCDKi to mdHIV #77 cells for 14 days after which the drug was washed (removed from the culture media) and cells were monitored for 10 more days by flow cytometry (Fig. 8*B*, 8*C*). YKL-5-124 (CDK7i) was not assessed given the toxicity of this drug when applied for several days (Fig. 6*A*, 6*B*).

**Figure 8.**
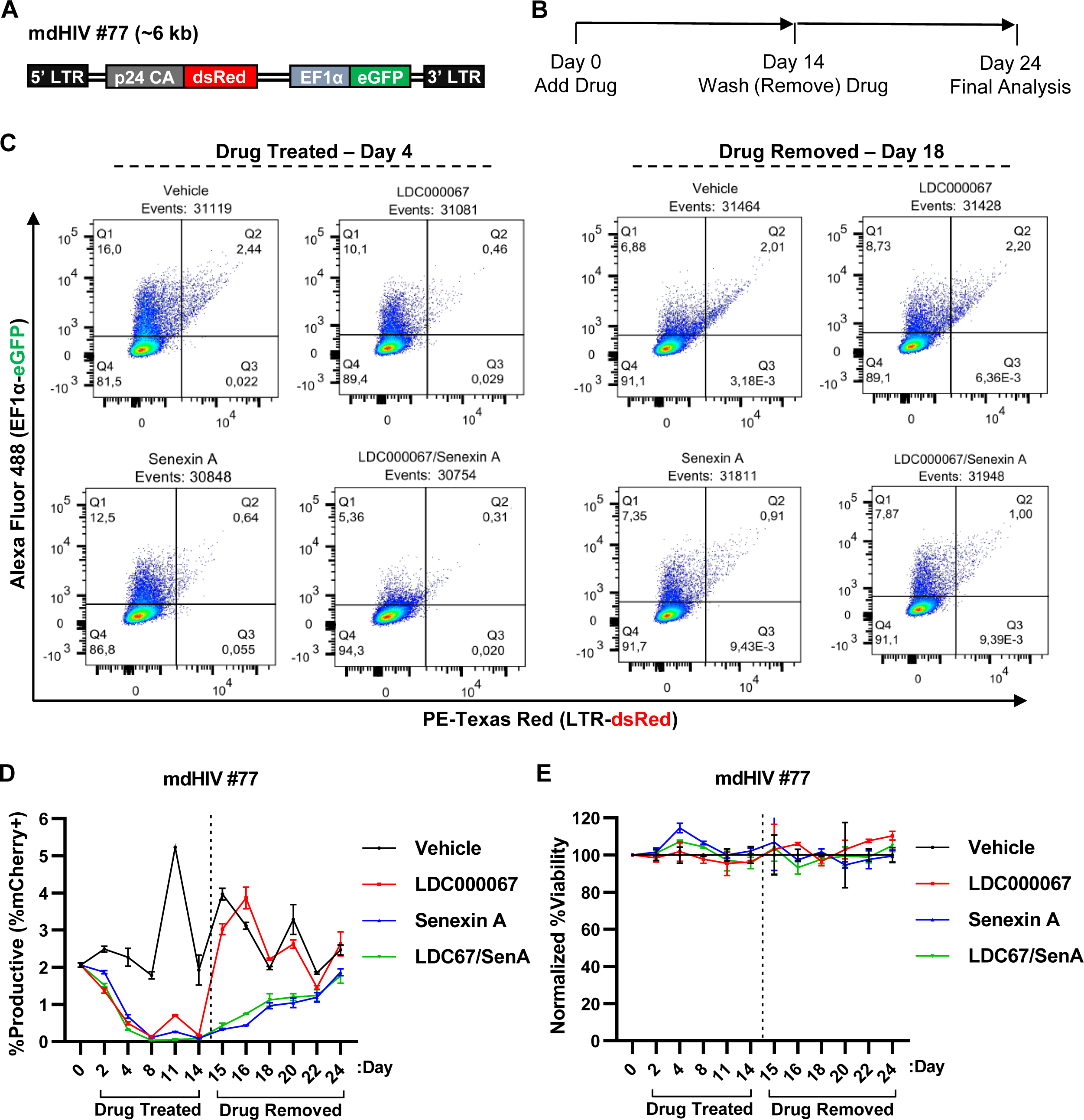
Effect of CDK9 and/or CDK8/19 inhibition on mdHIV #77 proviral latency. **A:** Simplified depiction of the replication incompetent mini-dual HIV-1 reporter virus that is chromosomally integrated in the mdHIV #77 Jurkat Tat clonal T cell line. 5’ LTR transcription drives dsRed expression as a fusion with p24 Gag while an internal EF1α promoter expresses eGFP. All other auxiliary proteins including Tat are not present. **B:** Experimental design. mdHIV #77 cells were incubated with a vehicle control (DMSO), 1 μM LDC000067, 10 μM Senexin A, or 1 μM LDC000067 and 10 μM Senexin A for 14 days after which the drug was removed from the media by washing and viral expression was assessed by flow cytometry for 10 more days. **C:** Representative flow cytometry scatter plots following 4 days of drug treatment and 4 days following drug withdrawal. All “dots” are indicative of a single HIV-1 infected cell with Q2 (dsRED+/eGFP+) possessing cells with transcriptionally active provirus. **D:** mdHIV #77 cells were treated as in (**B**) with viral expression assessed on the indicated day by flow cytometry (*n* = 2, mean ±LSD). **E:** mdHIV #77 cells treated as indicated in (**B**) were examined for viability, with values normalized to the vehicle control (*n* = 2, mean ±LSD).

Consistent with our findings that tCDK activity is required for reactivation of latent provirus (Fig. 2) and stimulates transcription early upon infection (Fig. 7), basal HIV-1 expression was practically outright suppressed by LDC000067 (CDK9i) and/or Senexin A (CDK8/19i) following 4 days of treatment (Fig. 8*C*, 8*D*). To great surprise, removal of the CDK9 inhibitor LDC000067 corresponded with immediate rebound of proviral expression as the population of dsRed+/eGFP+ cells nearly matched that of uninhibited cells within 24 hrs of drug withdrawal (Fig. 8*D*, compare day 14 to 15). Very importantly however, mdHIV #77 cells that had been treated with Senexin A (CDK8/19i) alone or in combination with LDC000067 (CDK9i) displayed maintenance of latency, with the tCDKi treated cells displaying reduced HIV-1 expression for more than 10 days following drug removal (Fig. 8*D*, Drug Washed). Of note, none of the inhibitor treatments affected cellular viability (Fig. 8*E*). Additionally, we examined the Jurkat Tat mdHIV #110 clonal line (Fig. S6). Akin to mdHIV #77 cells, the mdHIV #110 cell line possesses a chromosomally integrated mini-dual reporter provirus (Fig. S6*A*), however, this virus is located in *BTBD10* in the (-) orientation and has a proportion of ∼25% productive cells (Hashemi *et al*, 2016). Treatment of mdHIV #110 resulted in similar observations as either inhibitor used alone or in combination reduced the population of productive provirus (Fig. S6*D*, Drug Treated). Once again, latency caused by LDC000067 (CDK9i) was not robust, with viral expression rebounding within 24 hrs of withdrawal (Fig. S6*D*, compare day 14 to 15). In contrast, Senexin A (CDK8/19i) treated cells maintained lower viral expression days following removal (Fig. S6*D*, Drug Washed). The viability of mdHIV #110 cells was unaffected by the tCDKi treatments (Fig. S6*E*).

Above, we examined the capability of CDK9 and/or CDK8/19 inhibitors to generate robust, lasting HIV-1 latency (Fig. 8, Fig. S6). However, these cell lines were generated from Jurkat T cells that constitutively express Tat and with a mini-HIV-1 reporter virus that lacks expression of viral proteins (Hashemi *et al*, 2016). As such, we wanted to confirm our findings using a full-length HIV-1 provirus with auxiliary protein expression, including Tat, coupled to 5’ LTR transcriptional activity. To this end, we generated a clonal RGH Jurkat T cell line (Fig. 9*A*, RGH#20) and treated it for 7 days with LDC000067 (CDK9i) and/or Senexin A (CDK8/19i) with analysis continuing for 10 days following drug withdrawal (Fig. 9*B*). Near complete viral latency was established following 4 days of inhibitor treatment (Fig. 9*C*, 9*D*, Drug Treated). Again, removal of the CDK9 inhibitor LDC000067 was followed by viral rebound while cells retained decreased HIV-1 transcription for more than a week following the removal of the CDK8/19 inhibitor Senexin A (Fig. 9*D*, Drug Washed). Furthermore, no change in RGH #20 cell viability was observed for any of the administered treatments (Fig. 9*E*). Collectively, these results indicate that inhibitors of CDK9 will be incapable of producing long-lasting robust latency and as such, are not ideal “block and lock” LPA candidates. In contrast, inhibition of the Mediator kinases CDK8/CDK19 generates proviral latency that is maintained following drug withdrawal making them attractive targets for “block and lock” therapeutic approaches.

**Figure 9.**
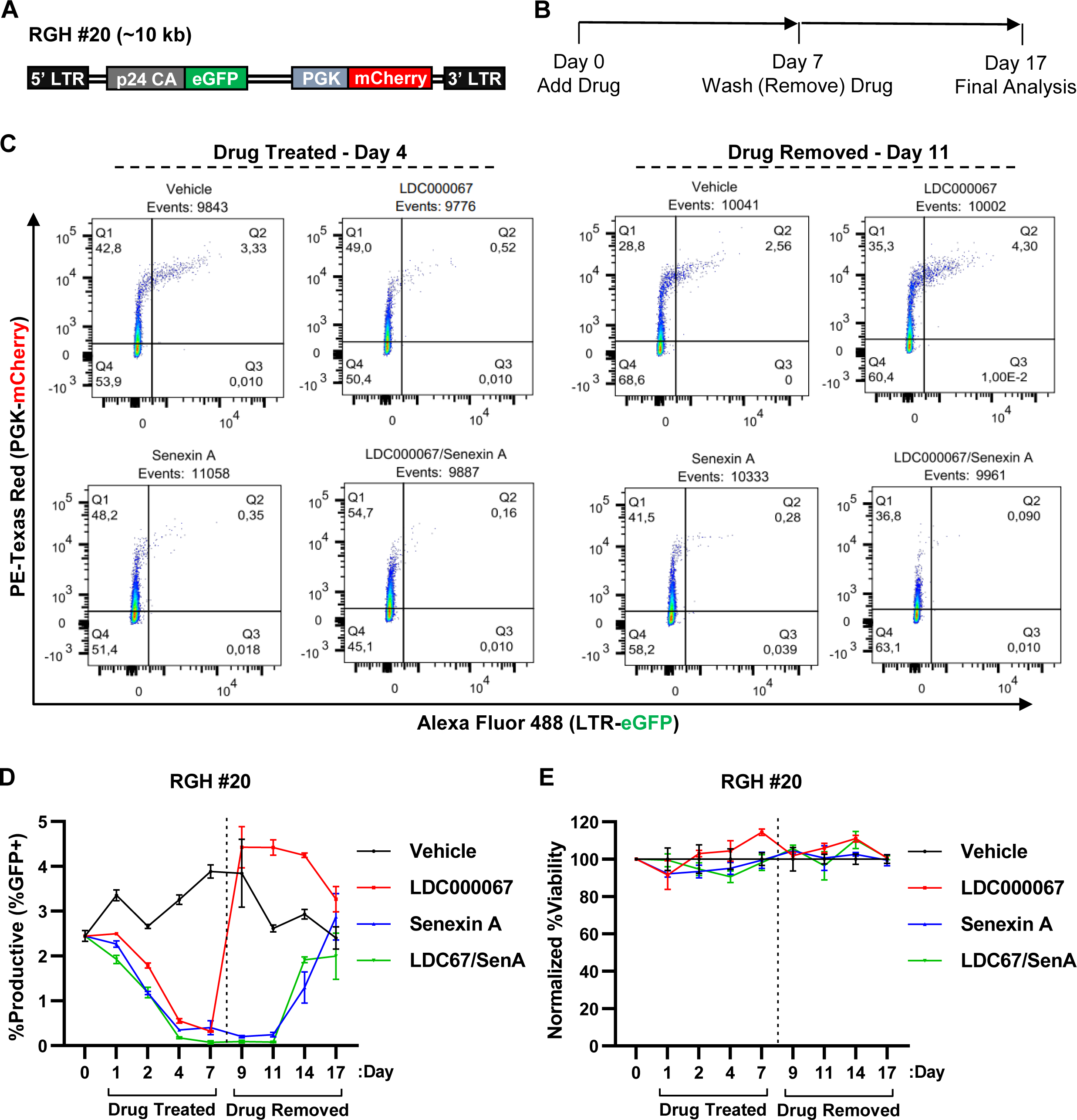
Effect of CDK9 and/or CDK8/19 inhibition on RGH #20 proviral latency. **A:** Simplified depiction of the full-length, replication incompetent HIV-1 reporter virus that is chromosomally integrated in the RGH #20 Jurkat clonal T cell line. 5’ LTR transcription drives eGFP expression as a fusion with p24 Gag while mCherry is expressed from an internal PGK promoter that replaces *Nef*. Other than *Nef*, all auxiliary proteins are expressed while a mutation in *Env* renders the provirus replication incompetent **B:** Experimental design. RGH #20 cells were incubated with a vehicle control (DMSO), 1 μM LDC000067, 10 μM Senexin A, or 1 μM LDC000067 and 10 μM Senexin A for 7 days after which the drug was removed from the media by washing and viral expression was assessed by flow cytometry for 10 more days. **C:** Representative flow cytometry scatter plots following 4 days of drug treatment and 4 days following drug withdrawal. All “dots” are indicative of a single HIV-1 infected cell with Q2 (eGFP+/mCherry+) displaying cells with transcriptionally active provirus. **D:** RGH #20 cells were treated as in (**B**) with viral expression assessed on the indicated day by flow cytometry (*n* = 2, mean ±LSD). **E:** RGH #20 cells treated as indicated in (**B**) were examined for viability, with values normalized to the vehicle control (*n* = 2, mean ±LSD).

### tCDK inhibitors suppress proviral expression in people living with HIV-1 ex vivo

We sought to confirm the requirement of tCDK activity for proviral expression by assessing the effect of kinase inhibition on primary CD4^+^ PBMCs isolated from individuals living with HIV-1 (Table 1). In all three of the individuals examined, we observed increased abundance in HIV-1 mRNA in response to PMA/ionomycin stimulation (Fig. 10, compare Unstimulated to Vehicle). In agreement with the results presented above, treatment with YKL-5-124 (CDK7i), LDC000067 (CDK9i), or Senexin A (CDK8/19i) consistently resulted in a decrease in proviral induction (Fig. 10). Interestingly, the combination of LDC000067 and Senexin A produced the most robust silencing of viral expression (Fig. 10, LDC000067/Senexin A), with Patient A displaying even lower quantities of HIV-1 mRNA than when left unstimulated (Fig. 10*A*). Furthermore, for Patient C we observe cooperative HIV-1 inhibition of combined LDC000067 and Senexin A treatment (Fig. 10*C*). Taken together, these results indicate that inhibition of the tCDKs can suppress reactivation of HIV-1 in patients *ex vivo*.

**Figure 10.**
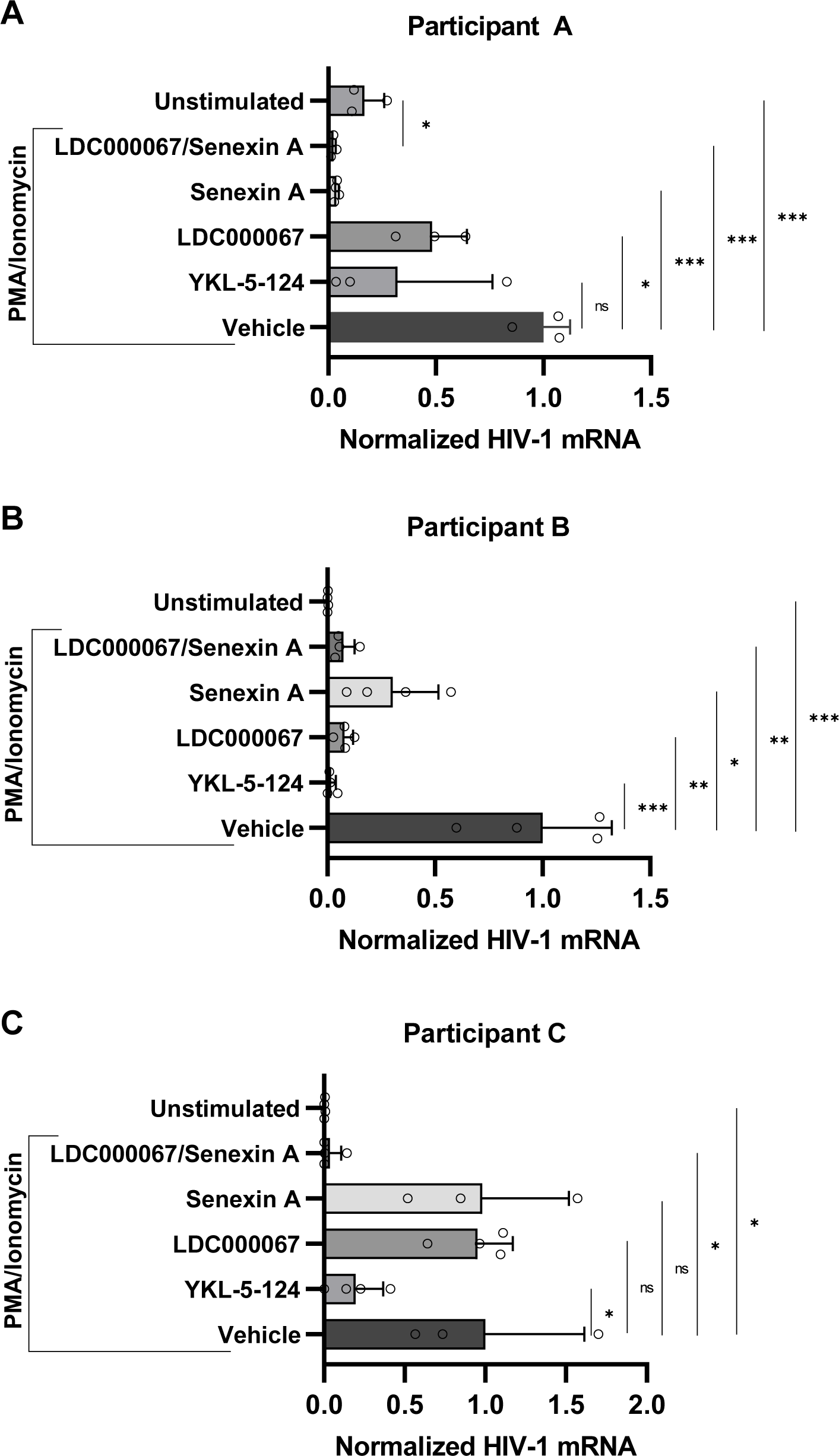
tCDK activity is required for reactivation of HIV-1 in patient CD4^+^ PBMCs *ex vivo*. **A - C:** CD4^+^ PBMCs, isolated from HIV-1 infected individuals on ART, were either left untreated or were pre-treated for 30 min with a Vehicle (DMSO), 100 nM YKL-5-124, 1 μM LDC000067, 10 μM Senexin A, or 1 μM LDC000067/ 10 μM Senexin A. Following the pre-treatment, cells were incubated with 5nM PMA/ 1μM Ionomycin for 24 hrs after which intracellular RNA was extracted and analyzed using RT-PCR with oligos specific for multiply spliced Tat-Rev HIV-1 mRNA transcripts. HIV-1 mRNA expression is normalized to *GAPDH* (*n* = 3 - 4, mean ±LSD).

**Table 1.**

| Patient | ID | Sex | Intact HIV Copies/Million<br>CD4+ T cells | Total HIV Copies/Million<br>CD4+ T cells | %Intact |
| --- | --- | --- | --- | --- | --- |
| A | BC024 | Male | 339.88 | 4941.32 | 6.88 |
| B | BC030 | Male | 397.27 | 3774.98 | 10.52 |
| C | BC002 | Male | 197.31 | 4857.65 | 4.06 |

## Discussion

Current therapeutic strategies to eliminate the life-long requirement of ART for people living with HIV-1 are aimed at modulating proviral latency. The “shock and kill” strategy of latency reversal has been the subject of intense investigation and although a number of LRAs have been identified, none have succeeded in reducing the pool of latently infected CD4^+^ T cells (Sadowski & Hashemi, 2019). The antithetical “block and lock” approach has been proposed to be a more realistic strategy to functionally eliminate HIV-1. The aforementioned therapeutic strategy relies on LPA mediated suppression of 5’ LTR transcription which may lead to sustained epigenetic silencing of integrated provirus (Moranguinho & Valente, 2020) (Vansant *et al*, 2020). Although several LPAs such as didehydro-cortistatin A (dCA) (Mousseau *et al*, 2015) (Kessing *et al*, 2017) and spironolactone (Mori *et al*, 2021) have displayed promising results, no human trial with any latency enforcing compound has been performed. In the present study, we assessed the transcriptional Cyclin Dependent Kinases (tCDKs) CDK7, CDK9, and CDK8/19 as potential targets for the robust enforcement of proviral latency in various cell line models and primary CD4^+^ T cells from people living with HIV-1.

Recruitment of the general transcription factor TFIIH to the LTR is a rate limiting step for reactivation of latent HIV-1 (Kim *et al*, 2006). TFIIH is a complex composed of 10 proteins consisting of a core component (XPB, XPD, p62, p52, p44, p34, p8) and a CDK-activating kinase (CAK) module (CDK7, cyclin H, MAT1) (Yan *et al*, 2019). In agreement with TFIIH recruitment being required for viral reactivation, we found that treatment with the selective CDK7 inhibitor YKL-5-124 prevented transcriptional induction of HIV-1 in response to T cell activation (Fig. 2). Furthermore, YKL-5-124 was found to limit reactivation in response to several diverse LRAs including the TRIM24 bromodomain inhibitor IACS-9571 and the HDACi SAHA (Fig. 4). Surprisingly, CDK7 inhibition with concomitant JQ1 mediated induction resulted in an increased proportion of cells possessing productive HIV-1 (Fig. 4*A*), although viral expression was severely dampened (Fig. 4*B*). Interestingly, we observed a similar expression pattern when we assessed the reactivation of YKL-5-124 treated JLatA72 cells (Fig. 5). The JLatA72 cell line possesses an integrated HIV-1 reporter that lacks Tat and as such, it would be reasonable to attribute this expression pattern to a Tat dependence. However, inhibition of CDK9 does not affect the reactivation of JLatA72 cells (-Tat) (Fig. 5*B*) but does impede reactivation of JLat10.6 cells in response to JQ1 (Fig. 4). As mentioned above, TFIIH is a multi-subunit complex with both helicase and kinase activities that facilitate promoter opening and phosphorylate the CTD of RNAPII at Ser-5. Our peculiar finding that CDK7 inhibition increases the pool of productive infections in certain conditions likely reflects the diverse functions of TFIIH for transcription circuit procession.

Additionally, we found that the CDK7 inhibitor YKL-5-124 was unable to prevent HIV-1 expression at concentrations that did not cause major disturbances in cell cycle progression (Fig. 6*A*, 6*B*). Perhaps even more intriguing, we observed that CDK7 inhibition was associated with increased susceptibility of T cells for HIV-1 infection (Fig. 7). Taken together, our assessment suggests that CDK7 inhibitors such as YKL-5-124 are unlikely to be appropriate for therapeutic use. Spironolactone causes the proteasomal degradation of the XBP subunit of TFIIH and has been shown to enforce HIV-1 latency (Mori *et al*, 2021). As such, spironolactone has been suggested as an LPA suitable for “block and lock” although viral rebound occurs immediately following drug withdrawal (Mori *et al*, 2021). Interestingly, although spironolactone severely restricts cellular XPB protein, it also leads to a reduction in CDK7 (Mori *et al*, 2021). Given our findings, inhibitors of TFIIH should be approached cautiously with regards to application in clinical settings.

P-TEFb, a heterodimer of Cyclin T1 and CDK9, is recruited to the HIV-1 LTR TAR element through interaction with Tat and facilitates escape of promoter proximal paused RNAPII, resulting in a strong positive feedback loop (Karn, 2011). We sought to interrupt the CDK9-Tat feedback loop and enforce latency by applying the selective inhibitor of CDK9 kinase activity, LDC000067 (Albert *et al*, 2014). Here, we found that LDC000067 suppressed proviral induction in various cell models (Fig. 2, Fig. 4) and in people living with HIV-1 *ex vivo* (Fig. 10). We also confirmed the Tat dependence of CDK9 for HIV-1 expression as LDC000067 had no appreciable effect in the absence of the viral trans-activator of transcription (Fig. 5, Fig. S4). Given the predominate role of CDK9 for HIV-1 transcription, the enzyme has been proposed as a potential therapeutic target (Canduri *et al*, 2008) (Ahlenstiel *et al*, 2020) and to this point, the CDK9 inhibitors Roscovitine, flavopiridol, and seliciclib have been shown to supress HIV-1 replication (Wang e*t al*, 2001) (Schang *et al*, 2002) (Biglione *et al*, 2007) (Schonhofer e*t al*, 2021). However, the aforementioned compounds are properly described as pan-CDK inhibitors as they are not selective and cause global inactivation of RNAPII transcription and cell cycle arrest (Blagosklonny, 2004) (Chao & Price, 2001). Of slight surprise, we found that the selective inhibition of CDK9 with LDC000067 had no effect on cell division and no apparent toxicity at concentrations effective at restricting proviral expression (Fig. 6*C*, 6*D*).

As LDC000067 administration was well tolerated, we were able to assess the capability of CDK9 inhibition to generate robust, lasting latency. In all T cell models examined, LDC000067 treatment generated proviral latency without affecting cell growth (Fig. 8, Fig. 9, Fig. S6, see Drug Treated). Disappointingly though, removal of the CDK9 inhibitor from media by washing corresponded with immediate HIV-1 rebound (Fig. 8, Fig. 9, Fig. S6, see Drug Washed). There is great interest in disrupting the Tat-CDK9 positive feedback loop in order to initiate “block and lock”, and several Tat targeting mechanisms have been identified including degradation of Tat by the long noncoding RNA NRON (Li *et al*, 2016) and introduction of the Tat mutant, Nullbasic (Jin *et al*, 2016) (Jin *et al*, 2019). Given that the predominate role of Tat for HIV-1 replication is the recruitment of P-TEFb (CDK9/Cyclin T1) to the 5’ LTR promoter and that we have observed that CDK9 inhibition is unable to generate durable latency, future studies should closely assess the ability of Tat inhibition to create long-lasting latency.

Recently, work from our lab has identified that CDK8 is a coactivator of HIV-1 expression and small molecule inhibition of CDK8/19 kinase activity restricts proviral reactivation (Horvath *et al*, 2023). Similarly, here we show that Senexin A inhibition of CDK8/19 suppresses viral induction in response to a diverse set of LRAs (Fig. 2, Fig. 4). The function of CDK8/19 is not dependent on Tat (Fig 5, Fig. S4), consistent with previous findings that CDK8/19 inhibition limits reactivation of ACH2 and U1 proviral latency cell models (Horvath *et al*, 2023). Additionally, we display a synergistic interaction between the CDK8/19 inhibitor Senexin A and inhibitors of either CDK7 or CDK9, suggesting distinct mechanism(s) of action. Intriguingly, CDK8/19 inhibition was effective beyond temporally acute treatment. Application of Senexin A at the point of infection alone caused a significant decrease in the proportion of transcriptionally active provirus in the days that followed (Fig. 7). Furthermore, inhibition of CDK8/19 with Senexin A was able to generate latency that was maintained for days following removal of the drug (Fig. 8, Fig. 9, Fig. S6). Collectively, our results indicate that the Mediator complex kinases, CDK8 and CDK19, are attractive targets for the development of a “block and lock” therapeutic strategy of sustained latency.

In the present study, we have evaluated the therapeutic potential of targeting CDK7, CDK9, and/or CDK8/19 for a functional HIV-1 cure. Inhibition of CDK7 presented several deficiencies including cellular toxicity and susceptibility to further HIV-1 infection. Although the CDK9 inhibitor LDC000067 was well tolerated and suppressed proviral expression, durable latency was not manifested. On the other hand, the CDK8/19 inhibitor Senexin A restricted HIV-1 expression at concentrations that did not affect cellular growth while also showing the ability to establish latency that was maintained following drug withdrawal. These results indicate that continued treatment with LDC000067 (CDK9i) and/or Senexin A (CDK8/19i) could be used therapeutically to deplete low-level viral replication in patients on ART. However, inhibition of CDK8/19 has the possibility of maintaining viral suppression following discontinuation of treatment.

## Materials and Methods

### Cell culture, virus culture, and lentiviral transduction

Jurkat E6-1, Jurkat Tat mHIV-Luciferase, JLat10.6, JLat9.2, JLatA72, and GXR-5 CEM cells were cultured in Roswell Park Memorial Institute 1640 (RPMI-1640) medium supplemented with 10% FBS, penicillin [100 units/ml], streptomycin [100 g/ml], and L-glutamine [2 mM]. HEK293T were cultured in Dulbecco’s modified Eagle’s (DMEM) medium supplemented with 10% fetal bovine serum (FBS), penicillin [100 units/ml], streptomycin [100 g/ml], and L-glutamine [2 mM]). All cell lines were incubated in a humidified 37°C and 5% CO_2_ atmosphere.

Peripheral Blood Mononuclear Cells (PBMC) from participants with HIV-1 on ART were isolated from whole blood by density gradient centrifugation using Lymphoprep^TM^ and SepMate^TM^ tubes (StemCell Technologies), and cryopreserved. Upon thawing, PBMCs were cultured in RPMI supplemented with 10% FBS, penicillin [100 units/ml], streptomycin [100 g/ml], and L-glutamine [2 mM]. Samples from participants were collected with written informed consent under a protocol jointly approved by the research ethics boards at Providence Health Care/UBC and Simon Fraser University (certificate H16-02474).

Vesicular stomatitis virus G (VSV-G) pseudotyped viral stocks were produced by co-transfecting HEK293T cells with a combination of viral molecular clone, psPAX, and VSV-G at a ratio of 8 μg: 4 μg: 2 μg. Transfections were performed with polyethylenimine (PEI) at a ratio of 6:1 (PEI:DNA) in Gibco Opti-MEM^tm^. Lentiviral infections were performed by plating 1×10^6^ cells in 24-well plates with media containing 8 μg/mL polybrene and the amount of viral supernatant to give the desired multiplicity of infection (M.O.I.) as indicated. Plates were subsequently spinoculated for 1.5 hrs at 1500 rpm.

### RT-PCR

RNA was extracted from cells using the RNeasy Kit (Qiagen) and analyzed with the Quant Studio 3 Real-Time PCR system (Applied Biosystems) using *Power* SYBR® Green RNA-to-CT™ 1-Step Kit (Thermo Fisher) as per the manufacturer’s instructions. RT-PCR data was normalized to GAPDH expression using the ΔΔCt method as previously described (Livak & Schmittgen, 2001). Cycling parameters were as follows: 48 °C, 30 min, 1x; 95 °C, 10 min, 1x; 95 °C, 15 sec, 60 °C, 1 min, 60x. Primers were as follows: HIV-1 mRNA (multiply spliced Tat-Rev transcripts), Fwd 5’ CTTAGGCATCTCCTATGGCAGGA, Rev 5’ GGATCTGTCTCTGTCTCTCTCTCCACC; GAPDH, Fwd 5’ TGCACCACCAACTGCTTAGC, Rev 5’ GGCATGGACTGTGGTCATGAG.

### Flow cytometry

Cells were treated as indicated in the figure legends. Human T cell lineages were suspended in PBS and a BD Biosciences LSRII-561 system was used for flow cytometry with threshold forward scatter (FSC) and side scatter (SSC) parameters being set so that a homogenous population of live cells was counted. FlowJo software (TreeStar) was used to analyze data and determine the indicated Mean Fluorescence Intensity (MFI).

### Quantitative analysis of tCDK inhibitor latency promoting agent interaction

Bliss independence modeling was used as previously described (Horvath *et al*, 2023) to statistically assess the inhibitory activity of combinatorial tCDK inhibitor treatment on HIV-1 reactivation. The equation *Fa*_xy, P_ = *Fa_x_* + *Fa_y_* – (*Fa_x_*)(*Fa_y_*) defines the Bliss independence model, in which *Fa*_xy, P_ is the predicted fraction affected by a combination of drug *x* and drug *y* that is derived from the experimentally observed fraction affected by drug *x* (*Fa_x_*) and drug *y* (*Fa_y_*) individually. Comparison of the predicted combinatorial affect (*Fa*_xy, P_) with the experimentally observed impact (*Fa _xy_*_,*O*_) was then performed: Δ*Fa _xy_*□=□*Fa _xy_*_,*O*_□−□*Fa _xy_*_,*P*_. If Δ*Fa _xy_* is greater than 0, the combination of drugs *x* and *y* exceed that of the predicted affect indicating that the drugs display synergistic interaction. If Δ*Fa _xy_* = 0, the drug combination follows the Bliss model for independent action. If Δ*Fa _xy_* is less than 0, the drug interaction is antagonistic as the observed effect of the drug combination is less than predicted.

### Statistics and reproducibility

All replicates are independent biological replicates and are presented as mean values withL± standard deviation shown by error bars. The number of times that an experiment was performed is indicated in the figure legends. P-values were determined by performing unpaired samples *t*-test with the use of GraphPad Prism 9.0.0. Statistical significance is indicated at \**P* < 0.05, \*\**P* < 0.01, or \*\*\**P* < 0.001, with n.s. denoting non-significant *P* ≥ 0.05.

## Data availability

All data supporting the findings of this study are available within the article or from the corresponding author upon reasonable request (I. Sadowski).

## Acknowledgments

We thank the laboratory staff at the BC Centre for Excellence in HIV/AIDS for processing PBMCs from study participants. This research was supported by program project grant F16-01210, from the Canadian Institutes of Health Research (CIHR), and F19-05392 Discovery Grant from the Natural Sciences and Engineering Research Council of Canada (NSERC) (to I.S.). PBMC collection from participants with HIV-1 was supported by CIHR project grant PJT-159625 (to ZLB). ZLB is supported by a Michael Smith Health Research BC Scholar Award.

## Author Contributions

Experimental design was conceived by Horvath R. M who also performed all experiments. Brumme Z.L. provided PBMC samples from individuals on ART. Horvath R. M. and Sadowski I. wrote the manuscript.

## Legends to Supplementary Figures

**Figure S1. Effect of tCDK inhibition on cellular viability. A – C:** JLat10.6 cells were pre-treaterd 30 min with the indicated concentration of YKL-5-124 (CDK7i) (**A**), LDC000067 (CDK9i) (**B**), or Senexin A (CDK8/19i) (**C**) and were subsequently stimulated with 5 nM PMA/ 1 μM ionomycin. Following 20 hrs, cell viability was assessed and is normalized to cells absent tCDK inhibitor (*n* = 2, mean ±LSD). **D – F:** As in (**A – C**), but JLat9.2 cells were assessed (*n* = 2, mean ±LSD).

**Figure S2. Effect of combined tCDK inhibitor treatment on cellular viability. A – C:** JLat10.6 cells were pre-treated for 30 min with 100 nM YKL-5-124 (CDK7i) and the indicated concentration of LDC000067 (CDK9i) (**A**) or Senxexin A (CDK8/19i) (**B**), or were pre-treated 30 min with 1 μM LDC000067 and the indicated concentration of Senexin A (**C**). Subsequently, cells were incubated with 5 nM PMA/ 1 μM ionomycin for 20 hrs. Following incubation, cell viability was determined and normalized to cells absent tCDK inhibitor (*n* = 2, mean ±LSD).

**Figure S3. IACS-9571 strongly synergizes with PEP005 to reactivate latent HIV-1.** JLat10.6 cells were left untreated, or were incubated with 5 nM PEP005, 20 μM IACS-9571, or 5 nM PEP005/ 20 μM IACS-9571. Following 20 hrs, cells were assessed for GFP expression by flow cytometry as shown in Fig. 4A. Calculation of synergy between the indicated treatments was performed using Bliss Independence Modeling. Data are presented as the difference between the predicted and the observed fractional HIV-1 expression response to the given drug combination (*n* = 2, mean ±LSD). See Materials and Methods for more details.

**Figure S4. Tat dependence of tCDK for proviral induction A:** Representative flow cytometry scatter plots of GXR-5 CEM cells (-Tat) that were pre-treated for 30 min with a vehicle control (DMSO), 100 nM YKL-5-124 (CDK7i), 1 μM LDC000067 (CDK9i), or 50 μM Senexin A (CDK8/19i) and subsequently incubated with 5 nM PMA and 1 μM ionomycin for 20 hrs. **B, C:** GXR-5 cells were left untreated or pre-treated for 30 min with a vehicle control (DMSO), 100 nM YKL-5-124, 1 μM LDC00067, or 50 μM Senexin A prior to being incubated with 5 nM PMA/ 1 μM ionomycin. Following 20 hrs, flow cytometry was performed with viral expression reported as the percent of GFP positive cells (**B**) and the delta GFP Mean Fluorescent Intensity (MFI) (**C**) (*n* = 2, mean ±LSD).

**Figure S5. Effect of tCDK inhibitors on cell viability during HIV-1 infection.** Jurkat human T cells were left untreated or were pre-treated for 30 min with 100 nM YKL-5-124, 1 μM LDC000067, or 10 μM Senexin A. Cells were subsequently infected with RG and 1-day post-infection, media was refreshed and cells were either washed of the drug or maintained with the same concentration of inhibitor as prior. 4 days post-infection, viability was determined and is normalized to untreated cells (*n* = 3, mean ±LSD).

**Figure S6. Effect of CDK9 and/or CDK8/19 inhibition on mdHIV #110 proviral latency. A:** Simplified depiction of the replication incompetent mini-dual HIV-1 reporter virus that is chromosomally integrated in the mdHIV #110 Jurkat Tat clonal T cell line. 5’ LTR transcription drives dsRed expression as a fusion with p24 Gag while an internal EF1α promoter expresses eGFP. All other auxiliary proteins including Tat are not present. **B:** Experimental design. mdHIV #110 cells were incubated with a vehicle control (DMSO), 1 μM LDC000067, 10 μM Senexin A, or 1 μM LDC000067 and 10 μM Senexin A for 14 days after which the drug was removed from the media by washing and viral expression was assessed by flow cytometry for 10 more days. **C:** Representative flow cytometry scatter plots following 4 days of drug treatment and 4 days following drug withdrawal. All “dots” are indicative of a single HIV-1 infected cell with Q2 (dsRED+/eGFP+) possessing cells with transcriptionally active provirus. **D:** mdHIV #110 cells were treated as in (**B**) with viral expression assessed on the indicated day by flow cytometry (*n* = 2, mean ±LSD). **E:** mdHIV #110 cells treated as indicated in (**B**) were examined for viability, with values normalized to the vehicle control (*n* = 2, mean ±LSD).

