## Supplementary Information for "Small molecule inhibitors of transcriptional Cyclin Dependent Kinases impose HIV-1 latency, presenting “block and lock” treatment strategies"

**Legends to Supplementary Figures**

**Figure S1. Effect of tCDK inhibition on cell viability. A – C:** JLat10.6 cells were pre-treated 30 minutes with the indicated concentration of YKL-5-124 (CDK7i) (**A**), LDC000067 (CDK9i) (**B**), or Senexin A (CDK8/19i) (**C**) and were subsequently stimulated with 5 nM PMA/ 1 μM ionomycin. Following 20 hours, cell viability was assessed and the results are normalized to untreated cells (*n* = 2, mean ± SD). **D – F:** As in (**A – C**), but JLat9.2 cells were assessed (*n* = 2, mean ± SD).

**Figure S2**. **Effect of tCDK inhibitor combinations on cell viability. A – C:** JLat10.6 cells were pre-treated for 30 minutes with 100 nM YKL-5-124 (CDK7i) and the indicated concentration of LDC000067 (CDK9i) (**A**) or Senxexin A (CDK8/19i) (**B**), or were pre-treated 30 minutes with 1 μM LDC000067 and the indicated concentration of Senexin A (**C**). Subsequently, cells were incubated with 5 nM PMA/ 1 μM ionomycin for 20 hours. Following incubation, cell viability was determined and normalized to untreated cells (*n* = 2, mean ± SD).

**Figure S3. IACS-9571 causes synergistic activation of latent HIV-1 in combination with PEP005.** JLat10.6 cells were left untreated, or incubated with 5 nM PEP005, 20 μM IACS-9571, or 5 nM PEP005/ 20 μM IACS-9571. GFP expression was measured 20 hours later by flow cytometry (Fig. 4A). Calculation of synergy between the indicated treatments was performed using Bliss Independence Modeling. Data are presented as the difference between the predicted and observed fractional HIV-1 expression response to the given drug combination (*n* = 2, mean ± SD). See Materials and Methods for details.

**Figure S4. Requirement of Tat for tCDK dependent HIV-1 expression. A:** Representative flow cytometry scatter plots of GXR-5 CEM cells (-Tat) pre-treated for 30 minutes with a vehicle control (DMSO), 100 nM YKL-5-124 (CDK7i), 1 µM LDC000067 (CDK9i), or 50 µM Senexin A (CDK8/19i) and subsequently incubated with 5 nM PMA and 1 µM ionomycin for 20 hours. **B, C:** GXR-5 cells were untreated or pre-treated for 30 minutes with a vehicle control (DMSO), 100 nM YKL-5-124, 1 µM LDC00067, or 50 µM Senexin A prior to incubation with 5 nM PMA/ 1 µM ionomycin. Following 20 hours, flow cytometry was performed with viral expression reported as the percent of GFP positive cells (**B**) and the delta GFP Mean Fluorescence Intensity (MFI) (**C**) (*n* = 2, mean ± SD).

**Figure S5. Effect of tCDK inhibitors on cell viability during HIV-1 infection.** Jurkat human T cells were left untreated or were pre-treated for 30 minutes with 100 nM YKL-5-124, 1 µM LDC000067, or 10 µM Senexin A. Cells were subsequently infected with RGH and 1-day post-infection, media was refreshed and cells were either washed of the drug or maintained with the same concentration of inhibitor. 4 days post-infection, viability was determined and is normalized to untreated cells (*n* = 3, mean ± SD).

**Figure S6. Effect of CDK9 and/or CDK8/19 inhibition on mdHIV #110 proviral latency. A:** Simplified depiction of the replication incompetent mini-dual HIV-1 reporter virus that is chromosomally integrated in the mdHIV #110 Jurkat Tat clonal T cell line (51), where dsRed is expressed from the 5' LTR as a fusion with p24 Gag while an internal EF1α promoter expresses eGFP. All other viral proteins including Tat are not expressed from the mini virus. **B:** mdHIV #110 cells were incubated with a vehicle control (DMSO), 1 μM LDC000067, 10 μM Senexin A, or 1 μM LDC000067 and 10 μM Senexin A for 14 days, after which the drug was removed from the media by washing and viral expression was assessed by flow cytometry for 10 more days. **C:** Representative flow cytometry scatter plots following 4 days of drug treatment and 4 days following drug withdrawal. All "dots" are indicative of a single HIV-1 infected cell with Q2 (dsRed+/eGFP+) possessing cells with transcriptionally active provirus. **D:** mdHIV #110 cells were treated as in (**B**) with viral expression assessed on the indicated day by flow cytometry ($n$ = 2, mean ± SD). **E:** mdHIV #110 cells treated as indicated in (**B**) were examined for viability, with values normalized to the vehicle control ($n$ = 2, mean ± SD).

### Figure S1

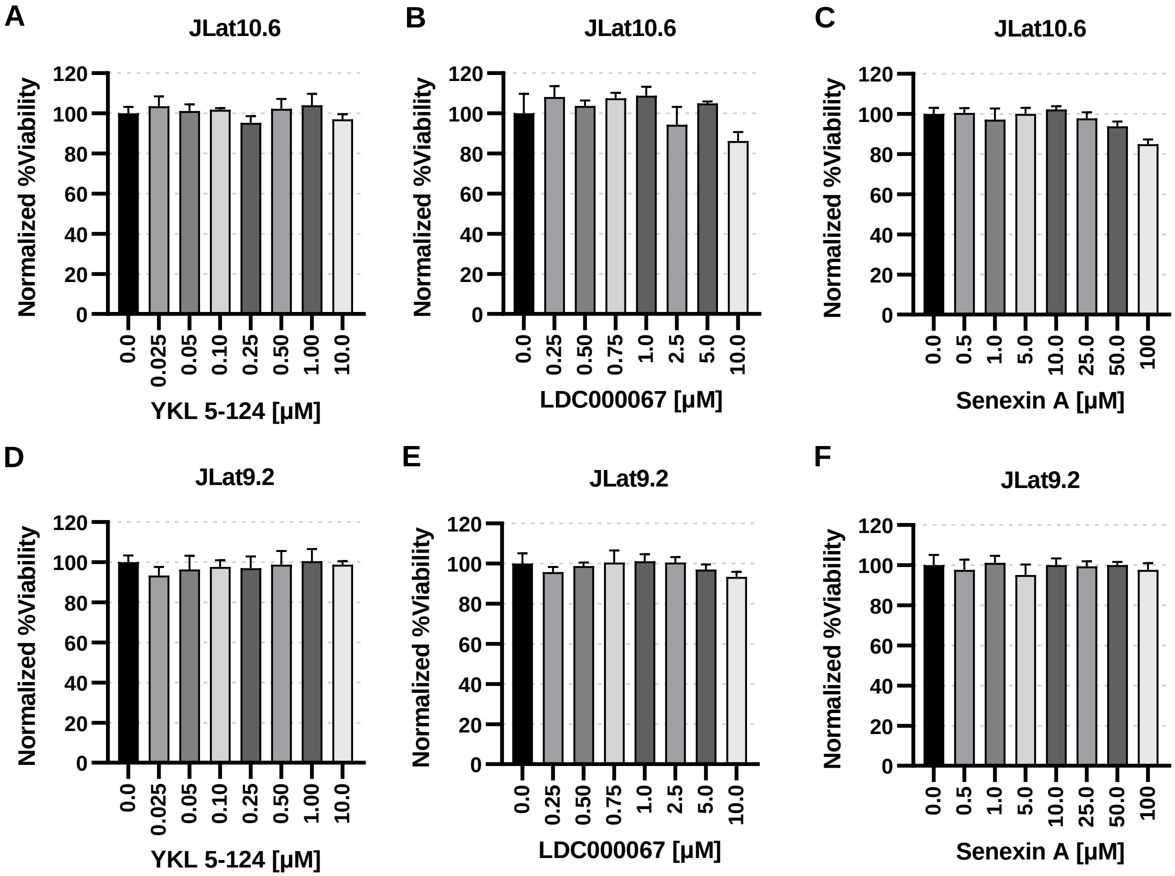
A
JLat10.6
Normalized %Viability
120
100
80
60
40
20
0
0.0
0.025
0.05
0.10
0.25
0.50
1.00
10.0
YKL 5-124 [μM]
B
JLat10.6
Normalized %Viability
0.0
0.25
0.50
0.75
1.0
2.5
5.0
10.0
LDC000067 [μM]
C
JLat10.6
Normalized %Viability
0.0
0.5
1.0
5.0
10.0
25.0
50.0
100
Senexin A [μM]
D
JLat9.2
Normalized %Viability
YKL 5-124 [μM]
E
JLat9.2
Normalized %Viability
LDC000067 [μM]
F
JLat9.2
Normalized %Viability
Senexin A [μM]

### Figure S2

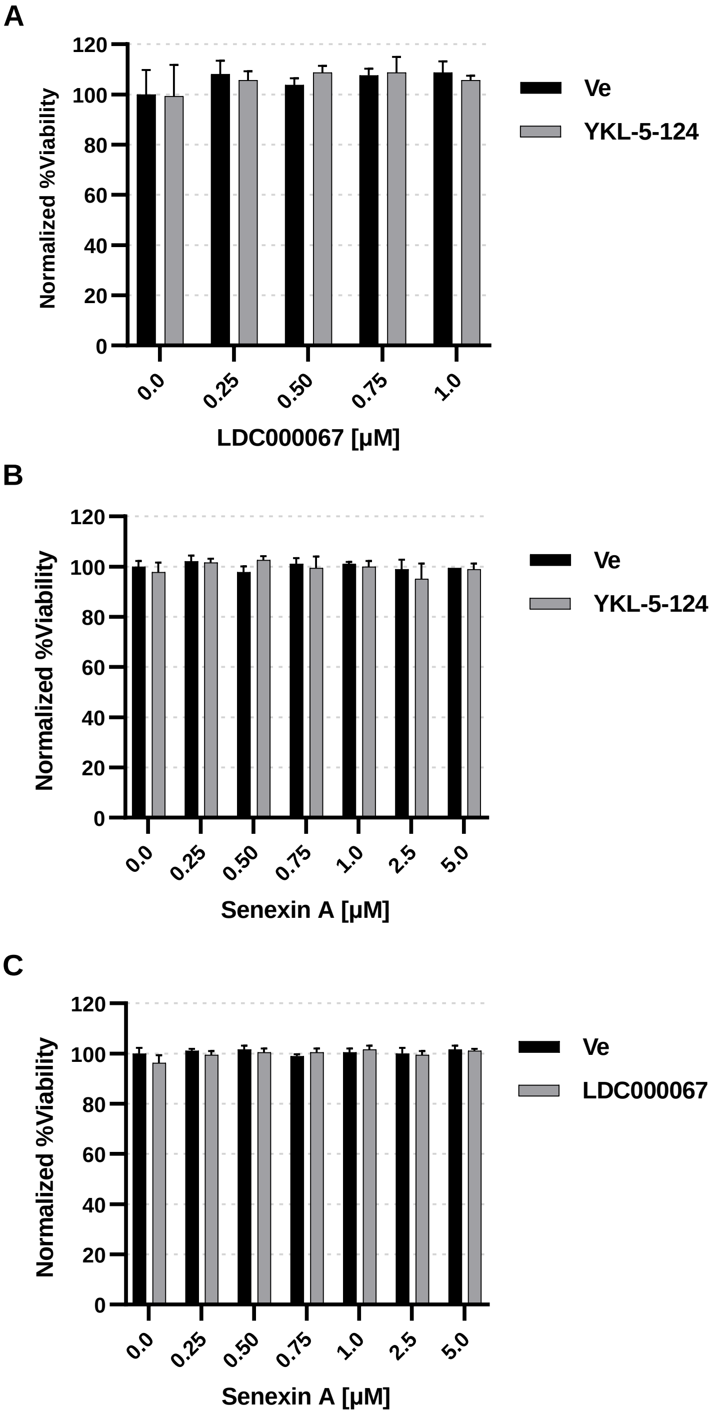
A
Normalized %Viability
120
100
80
60
40
20
0
0.0
0.25
0.50
0.75
1.0
LDC000067 [μM]
Ve
YKL-5-124
B
Normalized %Viability
120
100
80
60
40
20
0
0.0
0.25
0.50
0.75
1.0
2.5
5.0
Senexin A [μM]
Ve
YKL-5-124
C
Normalized %Viability
120
100
80
60
40
20
0
0.0
0.25
0.50
0.75
1.0
2.5
5.0
Senexin A [μM]
Ve
LDC000067

### Figure S3

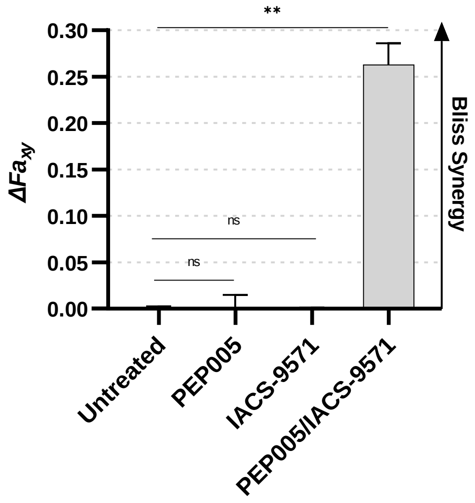

**
0.30
0.25
0.20
0.15
0.10
0.05
0.00
ΔFa_xy
ns
ns
Bliss Synergy
Untreated
PEP005
IACS-9571
PEP005/IACS-9571

### Figure S4

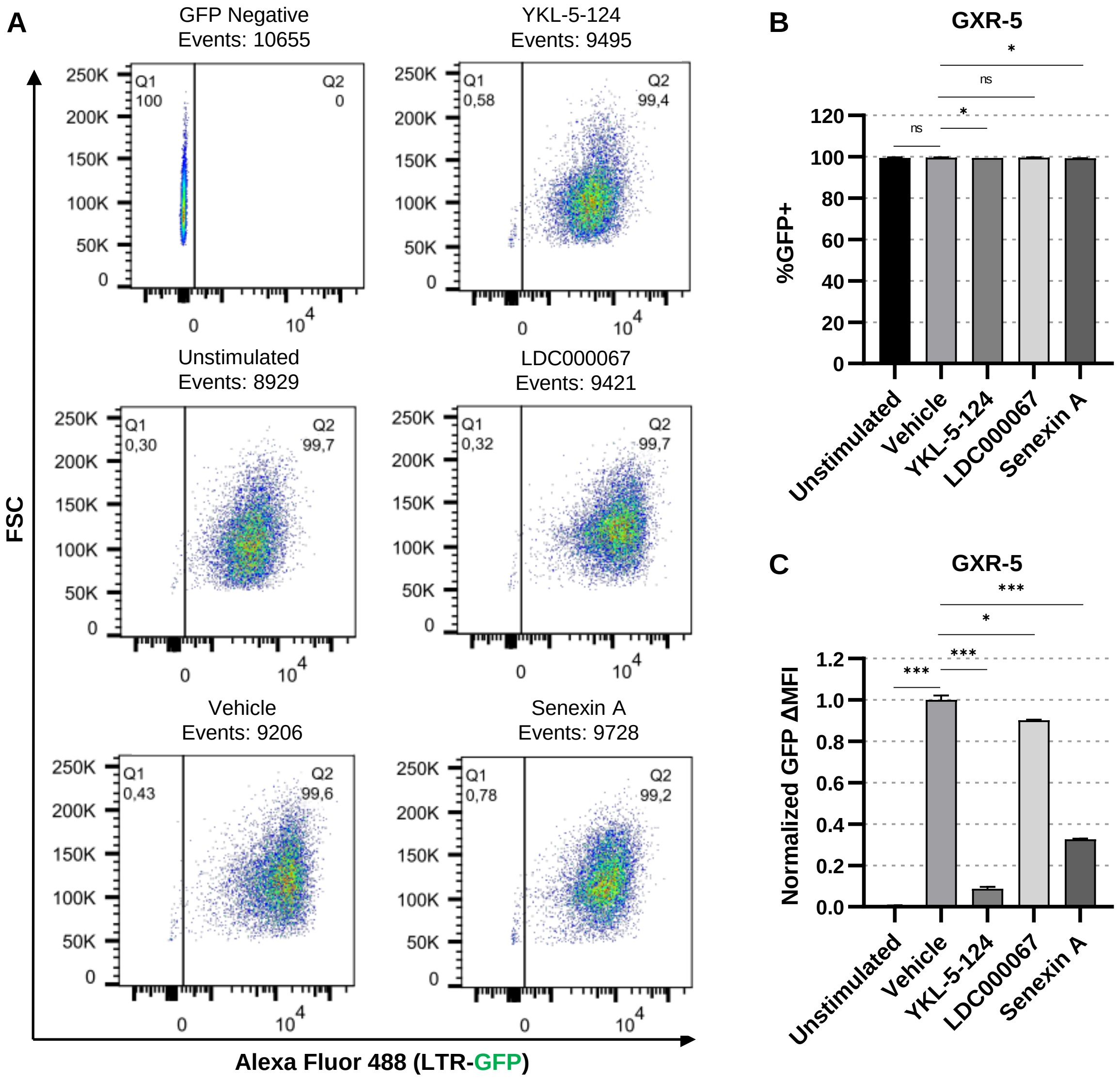

A
GFP Negative
Events: 10655
Q1
100
Q2
0
YKL-5-124
Events: 9495
Q1
0,58
Q2
99,4
Unstimulated
Events: 8929
Q1
0,30
Q2
99,7
LDC000067
Events: 9421
Q1
0,32
Q2
99,7
Vehicle
Events: 9206
Q1
0,43
Q2
99,6
Senexin A
Events: 9728
Q1
0,78
Q2
99,2
250K
200K
150K
100K
50K
0
0
10^4
FSC
Alexa Fluor 488 (LTR-GFP)
B
GXR-5
%GFP+
120
100
80
60
40
20
0
ns
*
ns
*
Unstimulated
Vehicle
YKL-5-124
LDC000067
Senexin A
C
GXR-5
Normalized GFP ΔMFI
1.2
1.0
0.8
0.6
0.4
0.2
0.0
***
***
*
***
Unstimulated
Vehicle
YKL-5-124
LDC000067
Senexin A

### Figure S5

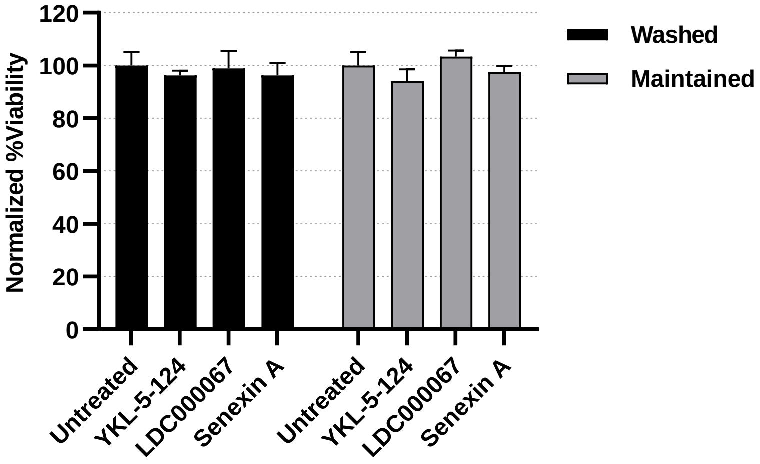

Normalized %Viability
120
100
80
60
40
20
0
Untreated
YKL-5-124
LDC000067
Senexin A
Untreated
YKL-5-124
LDC000067
Senexin A
Washed
Maintained

### Figure S6

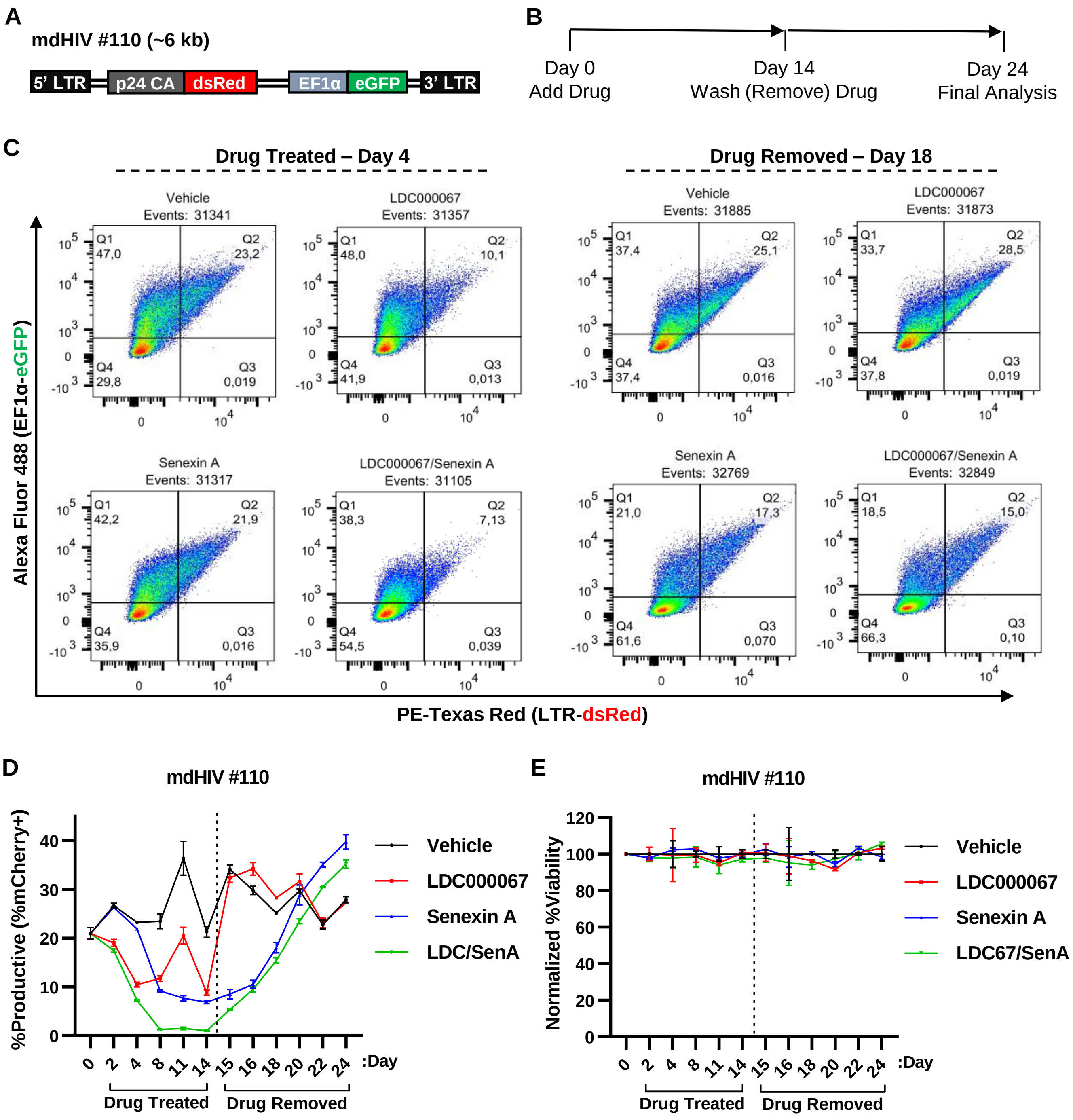
A
mdHIV #110 (~6 kb)
5' LTR
p24 CA
dsRed
EF1α
eGFP
3' LTR
B
Day 0
Add Drug
Day 14
Wash (Remove) Drug
Day 24
Final Analysis
C
Drug Treated – Day 4
Drug Removed – Day 18
Vehicle
Events: 31341
Q1 47,0
Q2 23,2
Q4 29,8
Q3 0,019
LDC000067
Events: 31357
Q1 48,0
Q2 10,1
Q4 41,9
Q3 0,013
Vehicle
Events: 31885
Q1 37,4
Q2 25,1
Q4 37,4
Q3 0,016
LDC000067
Events: 31873
Q1 33,7
Q2 28,5
Q4 37,8
Q3 0,019
Senexin A
Events: 31317
Q1 42,2
Q2 21,9
Q4 35,9
Q3 0,016
LDC000067/Senexin A
Events: 31105
Q1 38,3
Q2 7,13
Q4 54,5
Q3 0,039
Senexin A
Events: 32769
Q1 21,0
Q2 17,3
Q4 61,6
Q3 0,070
LDC000067/Senexin A
Events: 32849
Q1 18,5
Q2 15,0
Q4 66,3
Q3 0,10
Alexa Fluor 488 (EF1α-eGFP)
PE-Texas Red (LTR-dsRed)
D
mdHIV #110
%Productive (%mCherry+)
Vehicle
LDC000067
Senexin A
LDC/SenA
:Day
Drug Treated
Drug Removed
E
mdHIV #110
Normalized %Viability
Vehicle
LDC000067
Senexin A
LDC67/SenA
:Day
Drug Treated
Drug Removed
